# The Ancient Origin and Dynamic Diversification of the Fungal Poly(ADP)-ribose Polymerase Protein Family

**DOI:** 10.64898/2026.07.25.740670

**Authors:** Shira Milo, Cecelia N. Murphy, Madison Newman, Daniel Norment, Houlin Yu, Shay Covo, Li-Jun Ma

## Abstract

Poly(ADP-ribose) polymerases (PARPs) catalyze ADP-ribosylation, a conserved post-translational modification involved in DNA repair, transcriptional regulation, and chromatin remodeling. Although extensively studied in animals, the evolution and diversification of PARPs across the fungal kingdom remain largely unexplored. Here, we present the first kingdom-wide comparative genomic analysis of PARP proteins across 534 fungal species spanning eight phyla. We identified two primary conserved fungal PARP protein types corresponding to human PARP1 and PARP6. Both exhibit highly dynamic evolutionary histories characterized by frequent independent gain and loss events. Ancestral state reconstruction supports the presence of PARP1 in the last common ancestor of fungi, whereas the PARP6-like family has undergone repeated lineage-specific gains and losses. Fungal PARP6-like proteins retain a compact PARP catalytic domain fused to a C-terminal E2 ubiquitin-conjugating domain, whereas the PARP1 family displays extensive structural diversification through domain shuffling and lineage-specific fusions associated with DNA metabolism, chromatin remodeling, and signal transduction. Reconstruction of ancestral catalytic motifs across fungi and other eukaryotes revealed convergent evolution of a non-canonical H-Y-Y catalytic triad, with multiple motif variants co-occurring within individual proteins, suggesting functional diversification. In the *Fusarium oxysporum* species complex, we identified a lineage-specific expansion of the PARP family, driven exclusively by accessory chromosomes. Genomes with expanded PARP1 repertoires exhibited elevated basal PARylation, increased resistance to DNA-damaging agents that induce single strand breaks, and DNA damage-induced expression of accessory *Parp* genes. These findings reveal fungal PARPs as evolutionarily dynamic proteins that likely contribute to genome stability, adaptation, and pathogenicity.

## INTRODUCTION

Poly(ADP-ribose) polymerases (PARPs) are a family of enzymes that catalyze the transfer of ADP-ribose units from nicotinamide adenine dinucleotide (NAD⁺) to target substrates (Langelier et al., 2018) as part of the ADP-ribosyl transferase (ART) superfamily. Based on their enzymatic activity, PARPs are broadly divided into two functional classes: mono(ADP-ribosyl) transferases (MARylating PARPs), which transfer a single ADP-ribose moiety, and poly(ADP-ribosyl) polymerases (PARylating PARPs), which generate linear or branched chains of poly(ADP-ribose) (PAR). These modifications can be added to a variety of substrates, including proteins, DNA, or RNA, or occur as auto-modifications of the PARP enzyme itself (Perina et al., 2014). Cellular PARylation is transient, as PAR chains are rapidly degraded by poly(ADP-ribose) glycohydrolase (PARG) (Meyer-Ficca et al., 2004). PARP proteins have been identified across a broad range of eukaryotic lineages, from protists to humans, including in filamentous fungi (Citarelli et al., 2010).

The PARylation mechanism is evolutionarily conserved and is involved in a wide range of cellular processes, including DNA repair, transcriptional regulation, metabolism, and virulence (Perina et al., 2014). The functional importance of PARPs in human biology is demonstrated by extensive research on the 17 PARP family members encoded in the human genome, which participate in diverse cellular pathways including DNA repair, transcriptional regulation, chromatin remodeling, telomere maintenance, and immune responses (Citarelli et al., 2010; Ray Chaudhuri & Nussenzweig, 2017). Notably, these 17 PARPs exhibit limited functional redundancy, suggesting distinct and specialized roles for each family member (Vyas et al., 2013). Among the human PARP family members, PARP1 is the most extensively studied, primarily due to its central role in DNA repair and genome stability. PARP1 functions as a DNA damage sensor and repair mediator, rapidly recognizing DNA single-strand breaks and recruiting key factors involved in the base excision repair (BER) pathway, and to a lesser extent, in nucleotide excision repair (NER) and double-strand break (DSB) repair (Ray Chaudhuri & Nussenzweig, 2017). This critical role in genome maintenance has been successfully exploited for the development of PARP inhibitors that are now clinically approved and widely used in cancer treatment (Kaur et al., 2022).

Based on sequence homology of the PARP catalytic domain, the eukaryotic PARP family can be classified into six major clades (Citarelli et al., 2010). Clade 1 comprises the classical DNA-binding PARP1-3 proteins, which are primarily involved in DNA repair and transcriptional regulation (Perina et al., 2014). Clade 2 includes plant-specific PARPs, predominantly associated with stress responses (Rissel & Peiter, 2019). Clade 3 contains the majority of mono (ADP-ribosyl) transferases, including PARP7 and PARP9-15, which display considerable domain diversity and are implicated in a wide range of cellular functions (Citarelli et al., 2010; Feijs et al., 2013; Guo et al., 2004; Kalisch et al., 2012; Kleine et al., 2008). Clade 4 includes the tankyrases, which are PARylating enzymes involved in processes such as proteasome assembly and telomere maintenance (Cho-Park & Steller, 2013; Hsiao & Smith, 2008). Clade 5 consists of PARP4, also known as vault PARP (vPARP), which is found in a limited number of metazoans and amoebozoans (Perina et al., 2014). Clade 6 includes PARP6, PARP8, and PARP16 homologs, which are predicted to function primarily as mono(ADP-ribosyl) transferases (Gibson rissel& Kraus, 2012). Citarelli et al. (2010) suggested that the last common ancestor of all Eukaryotes encoded at least two types of PARP enzymes, a PARP1 and a PARP6 homolog. This hypothesis was later expanded to five different PARP enzymes presumably encoded by the last common ancestor of Eukarya (Perina et al., 2014).

The ART superfamily is primarily defined by conserved catalytic motifs that are essential for NAD⁺ binding and enzymatic activity. Two major families have been identified based on the conservation of three key residues: histidine-tyrosine-glutamate (H-Y-E), characteristic of the diphtheria toxin (DTX) family, and arginine-serine-glutamate (R-S-E), characteristic of the cholera toxin (CTX) family – both originally described in bacterial ARTs (Mikolčević et al., 2021). Within the PARP family, conservation of these catalytic residues varies among the six evolutionary clades. The canonical H-Y-E catalytic triad is retained in most members of Clades 1, 4, and 5, whereas substitutions of the catalytic glutamate are common in Clades 2 and 3. Clade 6 exhibits substitutions affecting both the first and third catalytic residues, resulting in multiple non-canonical catalytic signatures (Citarelli et al., 2010).

Fungi are a vital component of our ecosystem, significantly impacting agricultural productivity and public health as symbionts and pathogens of both plant and animal hosts (Rokas, 2022). The evolutionary success of fungi in colonizing a wide range of ecological niches has been linked to their remarkable genomic plasticity (Baroncelli et al., 2024; Sauters & Rokas, 2025; Selmecki et al., 2010; Vlaardingerbroek et al., 2016; Zhang et al., 2020) and the expansion of specific gene families associated with stress response, metabolism, and host adaptation (DeIulio et al., 2018; Yu et al., 2023). Among these, enzymes homologous to human PARP1 and PARP6 have been previously identified in fungal genomes (Citarelli et al., 2010; Perina et al., 2014). While several fungal PARP1 homologs have been functionally characterized, particularly in the context of growth, stress response (Kothe et al., 2010; Semighini et al., 2006) and, recently, virulence (Gao et al., 2024; J. Wang et al., 2024, 2025), their broader evolutionary significance and potential roles in fungal adaptation remain largely unexplored. In this study, we employ a comparative genomics and molecular evolution approach, taking advantage of the growing number of available fungal genome assemblies, to systematically analyze PARP proteins across the fungal kingdom. Our findings reveal the evolutionary trajectories of PARP enzymes across major fungal lineages and evaluate their potential contribution to the ecological and pathogenic diversity of fungi.

## RESULTS

### Highly dynamic and Ancient Origin of the Fungal PARP Proteins

Two primary types of PARP proteins were identified in fungi. One is a poly(ADP-ribosyl) polymerase human PARP1 homolog (hPARP1), characterized by a PARP catalytic domain, WGR domain, and PARP regulatory unit. The other is a mono(ADP-ribosyl) polymerase human PARP6- like homolog (hPARP6), containing a relatively short catalytic domain and a C-terminal E2 ubiquitin-conjugating (UBCc) domain, referred to as “PARP-Ubc” (Citarelli et al., 2010; Perina et al., 2014). To reach a comprehensive understanding of the conservation and evolutionary dynamics of fungal PARP proteins, we searched 534 fungal species representing eight major fungal phyla: Ascomycota, Basidiomycota, Blastocladiomycota, Chytridiomycota, Cryptomycota, Microsporidia, Mucoromycota, and Zoopagomycota, using a reciprocal best BLAST hit (rBBH) approach (Milo et al., 2019). The full-length PARP-Ubc sequence (*A. nidulans* XP_660733.1) and two PARP1 sequences (*A. nidulans* XP_660733.1, and Fol4287 XP_018243160.1) were used as queries. A total of 333 PARP1 orthologs were detected in six (Ascomycota, Basidiomycota, Chytridiomycota, Microsporidia, Mucoromycota and Zoopagomycota), while 322 PARP-Ubc were detected in four (Ascomycota, Basidiomycota, Chytridiomycota and Mucoromycota), out of eight phyla included in our dataset (Figure 1, Table S1-S2). Two of the phyla – Blastocladiomycota (single genome) and Cryptomycota/Rozellomycota (two genomes) – lack either form, although it may be due to the limited available genomes. We calculated the ancestral state and inferred gain or loss for each PARP protein by GLOOME (Cohen et al., 2010; Cohen & Pupko, 2011) using a stochastic mapping approach. Among 534 genomes, PARP1 and PARP-Ubc homologs are detected in 62% (333/534) and 61% (322/534) of the genomes, respectively. The GLOOME dynamics score, the sum of both gain and loss events, is 56.26 and 51.17 for PARP1 and PARP- Ubc, respectively, suggesting frequent and comparable gains and losses of both PARP proteins within the fungal kingdom.

**Figure 1.**
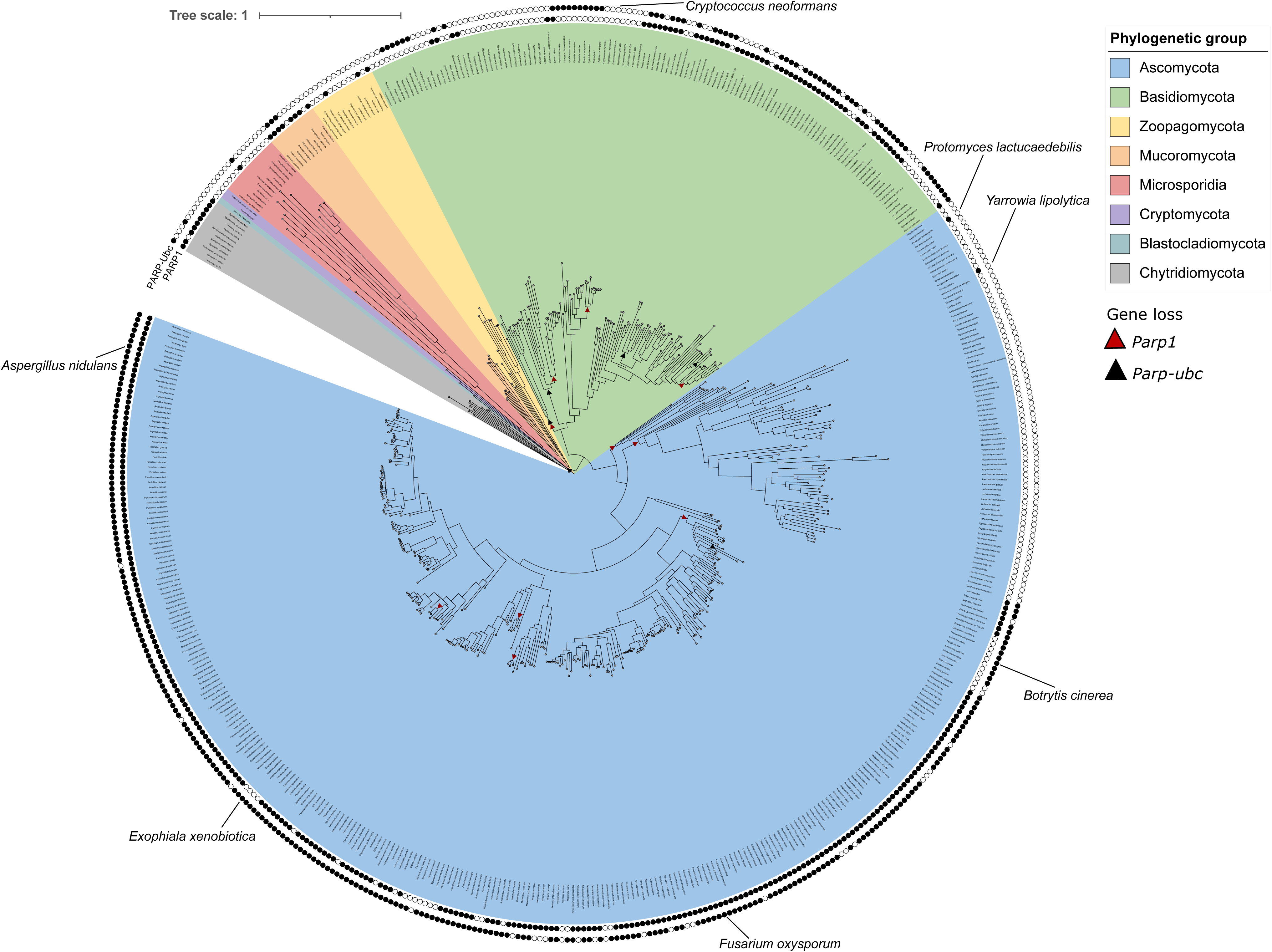
Conservation of PARP1 and PARP6 homologs within the fungal kingdom PARP1 and PARP-Ubc orthologs were searched using a reciprocal best BLAST hit (rBBH) approach. Fol4287 PARP1 as well as *A. nidulans* PrpA full protein sequences were used to identify orthologs across 534 fungal species from eight major phyla (Ascomycota, Basidiomycota, Blastocladiomycota, Chytridiomycota, Cryptomycota, Microsporidia, Mucoromycota, and Zoopagomycota). A genome-scale ML tree generated by (Li et al., 2021) was pruned, visualized and annotated with conservation state of the proteins; inner perimeter shows present/absent for PARP1 while outer perimeter shows present/absent for PARP-Ubc. Black circle indicates “present” whereas empty circle indicates “absent”. Loss events of PARP1 and PARP-Ubc inferred by GLOOME are indicated by red and black triangles, respectively.

Overall, there are fewer genomes available for early-diverging fungal lineages. Among four phyla, Microsporidia, Chytridiomycota, Zoopagomycota, Mucoromycota, we obtained 50 genomes, half of which (25) have both enzymes, while an additional 38% retained only one, often PARP1. Only one of the 12 Microsporidians genomes encodes PARP1, and none encode PARP- Ubc (Figure 1, Table S2). We observed a high retention rate of PARP1 among Chytrids, 10 out of 11 species, but low conservation of PARP-Ubc, presenting in two out of 11 species (Figure 1, Table S2). In the Zoopagomycota phylum, PARP-Ubc is absent and PARP1 homologs are detected in three out of 13 species (0.28% conservation rate) with multiple independent PARP1 loss events (Figure 1, Table S2). In the Mucoromycota phylum, eight out of 10 genomes retain PARP1 and three encode PARP-Ubc (Figure 1, Table S2). Three genomes within the subphylum Glomeromycotina retain both enzymes.

Among the 118 Basidiomycete fungal genomes, 56 (47%) and 62 (53%) genomes encode PARP1 and PARP-Ubc homologs, respectively (Figure 1, Table S2). Five classes (Malasseziomycetes, Mixiomycetes, Pucciniomycetes, Ustilaginomycetes, and Wallemiomycetes) completely lack both enzymes (Figure 1, Table S2) and GLOOME predicted early loss events on branches leading to Mixiomycetes and Pucciniomycetes. Tremellomycetes retained both forms, while PARP1 was lost in Cryptococcaceae, and a possible PARP-Ubc gain was predicted in the Microbotryomycetes class (Figure 1, Table S2). Agaricomycetes, with the largest available sequenced genomes in Basidiomycota, exhibited lineage-specific loss events, notably in the families of Agaricineae, Pluteineae, Russulales, and Hymenochaetaceae (Figure 1, Table S2).

A total of 254 and 256 (∼70%) of the 365 genomes within the Ascomycota phylum, encode PARP1 and PARP-Ubc homologs, respectively. Only two out of 78 (2.5%) yeast genomes (Pneumocystidomycetes, Saccharomycetes, Schizosaccharomycetes, Taphrinomycetes and Taphrinomycotina incertae sedis) harbor a PARP1, while none encode a PARP-Ubc homolog (Figure 1, Table S2). Two major, independent loss events of both enzymes were predicted on the branches leading to the subdivisions Taphrinomycotina and Saccharomycotina (Figure 1, Table S2). In both clades, GLOOME identified a single gain event of PARP1: in *Protomyces lactucaedebilis* (Taphrinomycotina) and *Yarrowia lipolytica* (Saccharomycotina) (Figure 1, Table S2). Among 288 filamentous Ascomycetes, 252 (87.5%) and 256 (88.8%) returned a positive hit for PARP1 and PARP-Ubc, respectively. Independent PARP1 loss events were predicted in Pezizomycotina: one within the class Leotiomycetes on the branch leading to Helotiales, followed by a gain in the Drepanopezizaceae, Mollisiaceae, and Ploettnerulaceae families. This was also accompanied by a loss of PARP-Ubc in the Erysiphales order (Figure 1, Table S2). Further PARP1 loss events were identified in the classes Dothideomycetes (Pseudocercospora genus) and Eurotiomycetes (Exophiala genus) (Figure 1, Table S2).

Together, these findings illustrate the evolutionary dynamics of PARP proteins in the fungal kingdom. The presence of a PARP1 homolog in the earliest diverging fungal clade Microsporidia, (Li et al., 2021) and the broad distribution of fungal PARP homologs support the hypothesis that the last common ancestor of fungi encoded a PARP1 homolog. The patchy distribution of PARP-Ubc among early-diverging fungi suggests a complex evolutionary history. Although ancestral-state reconstruction inferred absence of PARP-Ubc in the Microsporidia- Chytridiomycota ancestor. The observation of reduced representation among early-diverging fungal lineages and higher conservation rates among Dikarya, the higher fungal lineages, likely reflects the functional contribution of PARP proteins to the expansion of fungal lineages. The highly dynamic evolution pattern, marked by frequent gain and loss events, also suggests the plasticity and non-essential nature of fungal PARP proteins.

### Structural diversity and lineage-specific domain architecture of fungal PARP proteins

PARP-Ubc is present in 322 of 534 (60.3%) fungal genomes (Table S2) and is absent from multiple early-diverging fungal phyla (Microsporidia, Zoopagomycota, Blastocladiomycota). Defined by a bipartite domain architecture comprising an N-terminal PARP6 catalytic domain and a C-terminal E2 ubiquitin-conjugating enzyme catalytic domain (UBCc; UBE2Q subfamily), PARP-Ubc represents a fungal-specific function of ADP-ribosylation under the regulation of the ubiquitin-proteasome machinery (Citarelli et al., 2010). The canonical PARP-UBCc architecture (∼1154 aa mean length; range 481-1,324 aa) accounts for 299 of 322 (92.9%) proteins with domain data. Eighteen identified PARP-Ubc proteins (5.6%) retain only the PARP catalytic domain without a detectable UBCc domain (Table S2, architecture: PARP), including both Chytridiomycota representatives (*Synchytrium endobioticum* and *Piromyces* sp. E2; 736 and 232 aa, respectively), which may reflect an ancestral pre-fusion state or independent truncation. Five lineage-specific non-canonical fusions were additionally identified, all exclusively in Ascomycota and described in Table S2.

In contrast, PARP1 homologs exhibited substantial variation in both protein length and domain architecture while retaining a highly conserved catalytic core unit. All 333 identified PARP1 orthologs encode a PARP catalytic domain (*parp_like*, cd01437). Based on their core domain organization, fungal PARP1 proteins could be classified into three major architectural groups (Figure 2, Table S2). The predominant architecture, hereafter referred to as the canonical fungal PARP1, exist in 65% of the proteins (n = 217/333) and comprised the BRCT, WGR and PARP catalytic domains (mean length 724 aa). A second, structurally simplified architecture lacking the BRCT domain, was detected in 29% of the proteins (n = 96/333) and was significantly shorter than canonical PARP1 proteins (mean 609 aa; Mann-Whitney U test, p = 1.3 × 10^−^¹⁸). Finally, a small subset of proteins (7%; n = 22/333) acquired diverse lineage-specific domains through independent domain fusion events. These unique proteins were significantly longer than canonical PARP1 proteins (mean 1382 aa; Mann-Whitney U test, p = 2.6 × 10^−^⁵), reflecting the contribution of the additional domains. Comparison across fungal phyla revealed two major patterns of the diversification in PARP1 architecture: variation in the presence of the BRCT domain and lineage-specific acquisition of additional domains. The simplified WGR-PARP architecture is prevalent in most early diverging fungal lineages and was the only detected architecture in Zoopagomycota (n = 4/4; mean length 520 aa) and in 95% of Basidiomycota proteins (n = 53/56; mean length 611 aa). Within Chytridiomycota, both canonical and simplified architectures were detected, with BRCT-containing proteins present in a subset of Chytridiomycetes as well as in all sampled Monoblepharidomycetes and Neocallimastigomycetes. The Microsporidian PARP1 ortholog (182 aa) retained only a short catalytic PARP domain. In contrast, the canonical BRCT-containing architecture is commonly found in Pezizomycotina, accounting for 81% of proteins (n = 207/254). Accordingly, canonical PARP1 was significantly more frequent in Pezizomycotina than in all other fungal phyla combined (χ² = 121.3, df = 1, p = 3.3 × 10^−^²⁸), and Pezizomycotina proteins were significantly longer than those of Basidiomycota (mean 706 vs. 611 aa; Mann-Whitney U test, p = 4.8 × 10⁻¹³). Mucoromycota represents an intermediate pattern, with simplified architecture in 75% of the proteins (n = 6/8). Unexpectedly, the most structurally divergent PARP1 architecture was identified in four Mucoromycota species belonging to the classes Mucoromycetes and Umbelopsidomycetes. These proteins (2,648-2,995 aa) encode four to six tandem Ankyrin repeat (ANKYR) arrays N-terminal to the conserved WGR-PARP catalytic module. This architecture closely resembles vertebrate Tankyrase 1/2 (TNKS1/2), in which multiple ankyrin repeat clusters mediate substrate recognition. Similar proteins were previously identified in Amoebozoa (*D. discoideum*), Opisthokonta (*C. elegans*) and Chromalveolata (*S. diclina*) (Perina et al., 2014). Interestingly, in two Mucoromycota species, this protein harbors an additional CHROMO domain that was additionally detected only in *Fusarium decemcellulare*, where it is fused to integrase (Integrase_H2C2, pfam17921) and reverse transcriptase (RNase_HI_RT_Ty3) domains, suggesting that this chromodomain may participate more broadly in the mobilization of fungal genomic elements (Novikova, 2009). Among the 22 fusion proteins identified, 19 distinct domain architectures were detected, 16 of which occurred in a single species, indicating that most fusion events arose independently rather than being inherited from a closely-related common ancestor. Within Ascomycota, fusion frequency increases progressively in later-diverging classes: from none in Leotiomycetes and Pezizomycetes to 4% in Eurotiomycetes, 7% in Sordariomycetes, and 10% in Dothideomycetes, suggesting that novel architectures have primarily accumulated during the diversification of filamentous Ascomycota. The functional categories of acquired domains include protein degradation, transcriptional regulation, protein-protein interaction, DNA topology, and metabolic regulation (Figure 2, Table S2).

**Figure 2.**
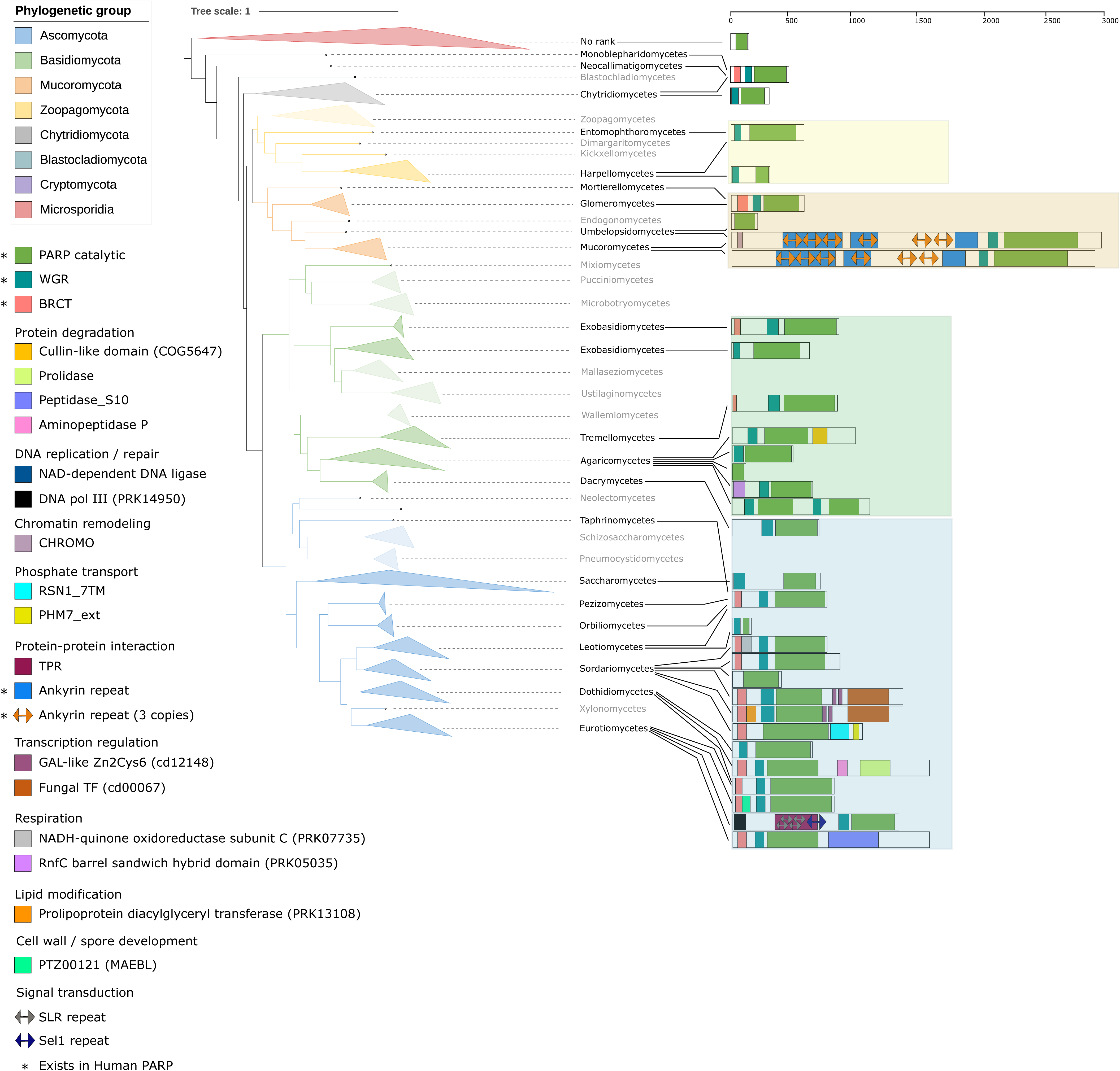
Domain architecture diversity of the fungal PARP1 protein PARP1 proteins of 534 fungal species from eight major phyla (Ascomycota, Basidiomycota, Blastocladiomycota, Chytridiomycota, Cryptomycota, Microsporidia, Mucoromycota, and Zoopagomycota) were subjected to conserve domain analysis using NCBI Batch CD-Search. A genome-scale Maximum likelihood tree generated by (Li et al., 2021) was pruned, visualized and annotated with representative PARP1 protein domain structures for each phylum/class. Colored branches indicate the phylogenetic placement of each species. Representative PARP1 domain architectures are grouped according to the major fungal lineages, indicated by shaded boxes. Protein domain/families that are found in human PARPs are marked with an asterisk.

Three cases are particularly informative regarding the origins and selective constraints that might be operating on these fusion events.

- Parylation through enhanced protein-protein interaction: In Eurotiomycetes, *Rasamsonia emersonii* (1332 aa) and *Talaromyces atroroseus* (1311 aa) encode PARP1 proteins carrying a C-terminal serine carboxypeptidase domain (Peptidase_S10) (Figure 2, Table S2). Independently, *Leptosphaeria maculans* (1667 aa) and *Pyrenochaeta* sp. (1558 aa) acquired C-

terminal M24 metallopeptidase modules (AMP_N–Prolidase complex) (Figure 2, Table S2), indicating repeated integration of proteolytic functions into PARP1. A structurally distinct architecture was identified in *Penicillium subrubescens*, where the N-terminal BRCT domain is replaced by extended TPR and Sel1 repeat arrays (Figure 2, Table S2). Although unrelated in sequence to ankyrin repeats, these interaction scaffolds may similarly expand substrate recognition through protein-protein interactions.

- PARylation and transcription regulation: A second recurrent pattern is the integration of transcriptional regulatory functions into the PARP1 protein in three closely related Sordariomycetes. *Thermothelomyces thermophila*, *Thielavia terrestris*, and *Chaetomium globosum*, encode PARP1 proteins fused to fungal-specific Zn₂Cys₆/GAL4 and fungal_TF_MHR transcription factor domains. *T. terrestris*additionally encodes an N-terminal LGT domain. This suggest that the functional association between PARylation and transcription regulation has been preferentially maintained within the lineage.
- A third pattern was observed in two *Colletotrichum* species, where distinct regulatory modules, a DTHCT domain (derived from DNA gyrase B/topoisomerase IV ATPase) in *C. sublineola* and an NADH-quinone oxidoreductase module in *C. incanum*, were independently inserted between the BRCT and WGR domains. The integration of functionally distinct domains in two closely related *Colletotrichum* species indicates that diversification of PARP1- associated cellular functions has occurred even over relatively short evolutionary distances.

Taken together, these results demonstrate that fungal PARP1 proteins exhibit extensive structural diversity across the fungal kingdom, ranging from a minimal catalytic form in early-diverging lineages to the canonical BRCT-containing architecture in Pezizomycotina, with additional lineage-specific domain fusions most pronounced in filamentous Ascomycota.

### Unique Expansion of PARP Proteins in the *Fusarium oxysporum* Species Complex

The analysis revealed increased structure and functional diversity in PARP1 proteins among Dikarya, and more so within Ascomycota. To further investigate PARP family dynamics, including family expansion and copy number variation, we analyzed 30 representative ascomycete genomes: two model yeasts (*S. cerevisiae*, *S. pombe*), four well-characterized filamentous fungi outside of the *Fusarium* genus (*A. acristatulus*, *A. nidulans*, *N. crassa*, *M. oryzae*), nine *Fusarium* species, and 15 strains from the *F. oxysporum* species complex (FOSC) (a complete list is found in Table S3). These taxa represent diverse phylogenetic relationships, lifestyles and ecological niches.

Using a conserved domain-based comparative genomics pipeline (see Methods), we defined PARP proteins as any containing a PARP catalytic domain based on InterProScan annotations (v5.38-76.0) (Blum et al., 2021) (Table 1, Table S4). As expected, the model yeasts lack detectable PARPs. Most filamentous Ascomycetes encode two to three PARP proteins, with *F. graminearum* harboring four (Figure 3A). Within the FOSC, however, PARP family size varied dramatically, ranging from two to twenty genes per genome (Figure 3B, Table 1). Across all non-FOSC Ascomycetes, PARP copy number is positively correlated with total proteome size (R² = 0.58; Figure 3C). In contrast, FOSC strains show no such correlation (R² = 0.11; Figure 3D), indicating that PARP family expansion in *F. oxysporum* (FO) is independent of whole-genome expansion that occurred during the evolution of Ascomycete fungi (Kelkar & Ochman, 2012; Milo et al., 2019). Furthermore, the number of PARP proteins does not correlate with specific lifestyle or host (Figure 3D). Interestingly, strains infecting Brassicaceae (f. sp. *conglutinans*, *raphani*, *matthiolae* PHW726, and Fo5176) encode the largest PARP repertoires (9, 9, 14, and 20 genes, respectively) and harbor significantly more PARP genes than plant-pathogenic FOSC strains infecting other hosts (mean 13.0 vs. 5.0; Mann-Whitney U test, p = 0.0051).

**Figure 3.**
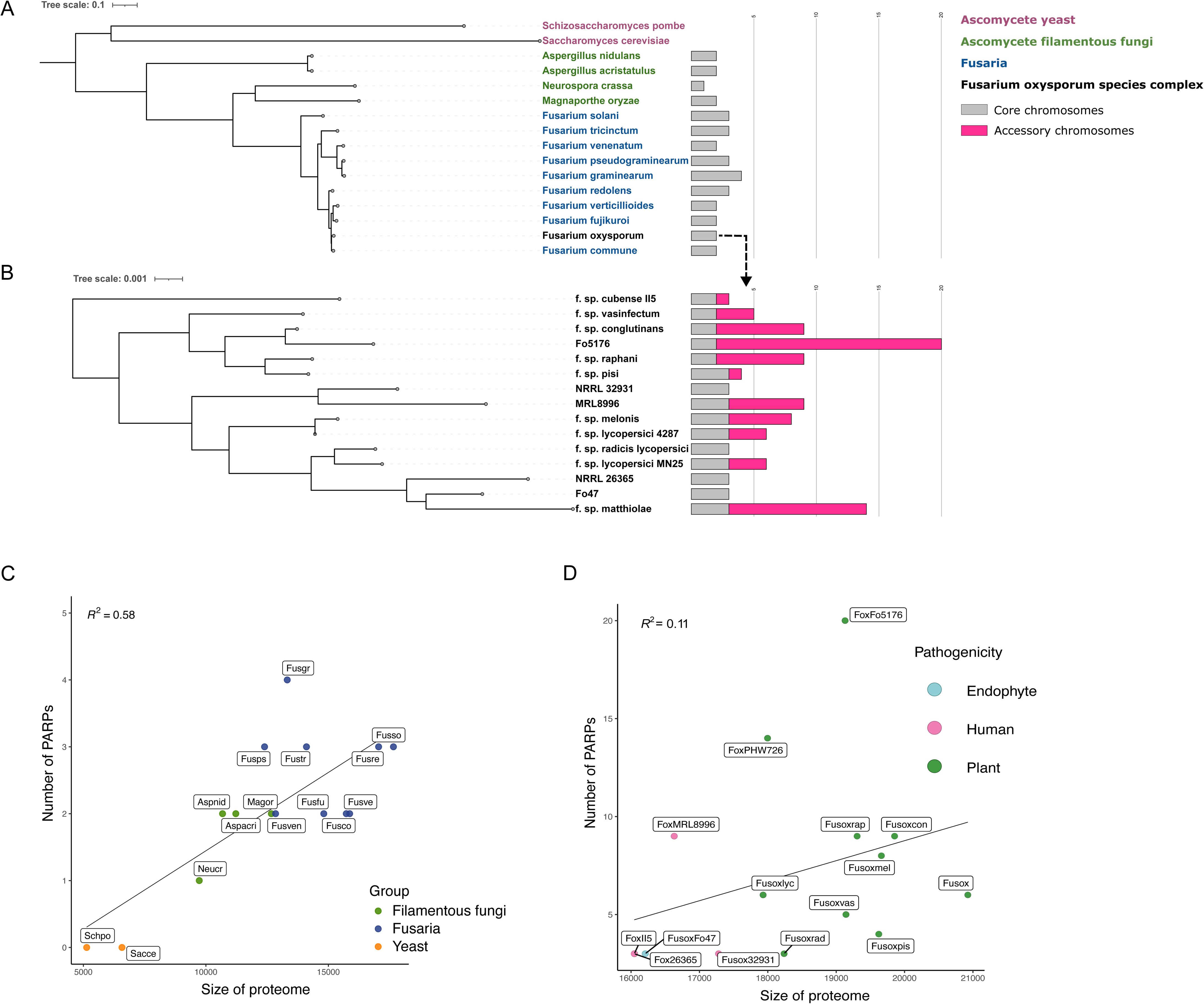
Lineage specific expansion of PARP proteins in the *F. oxysporum* species complex Phylogenetic tree of A) 15 Ascomycete species from three phylogenetic groups (yeasts, filamentous fungi, and Fusaria), and B) 15 FO strains, were constructed using a concatenated alignment of randomly selected 500 single-copy orthologous proteins to generate a Maximum likelihood tree of 1000 bootstraps. Trees were annotated with the *Parp* gene copy number in each genome. Gray bars show genes located on the core genome, whereas pink bars show genes located on accessory chromosomes. C) Correlation between the number of PARPs and proteome size in 15 Ascomycete species outside of the FOSC. Orange indicates yeasts, green indicates filamentous fungi, and blue indicates Fusaria. D) Correlation between the number of PARPs and proteome size in the FOSC. Green indicates plant pathogens, blue indicates endophytes, and pink indicates human pathogenic strains.

**Table 1.**
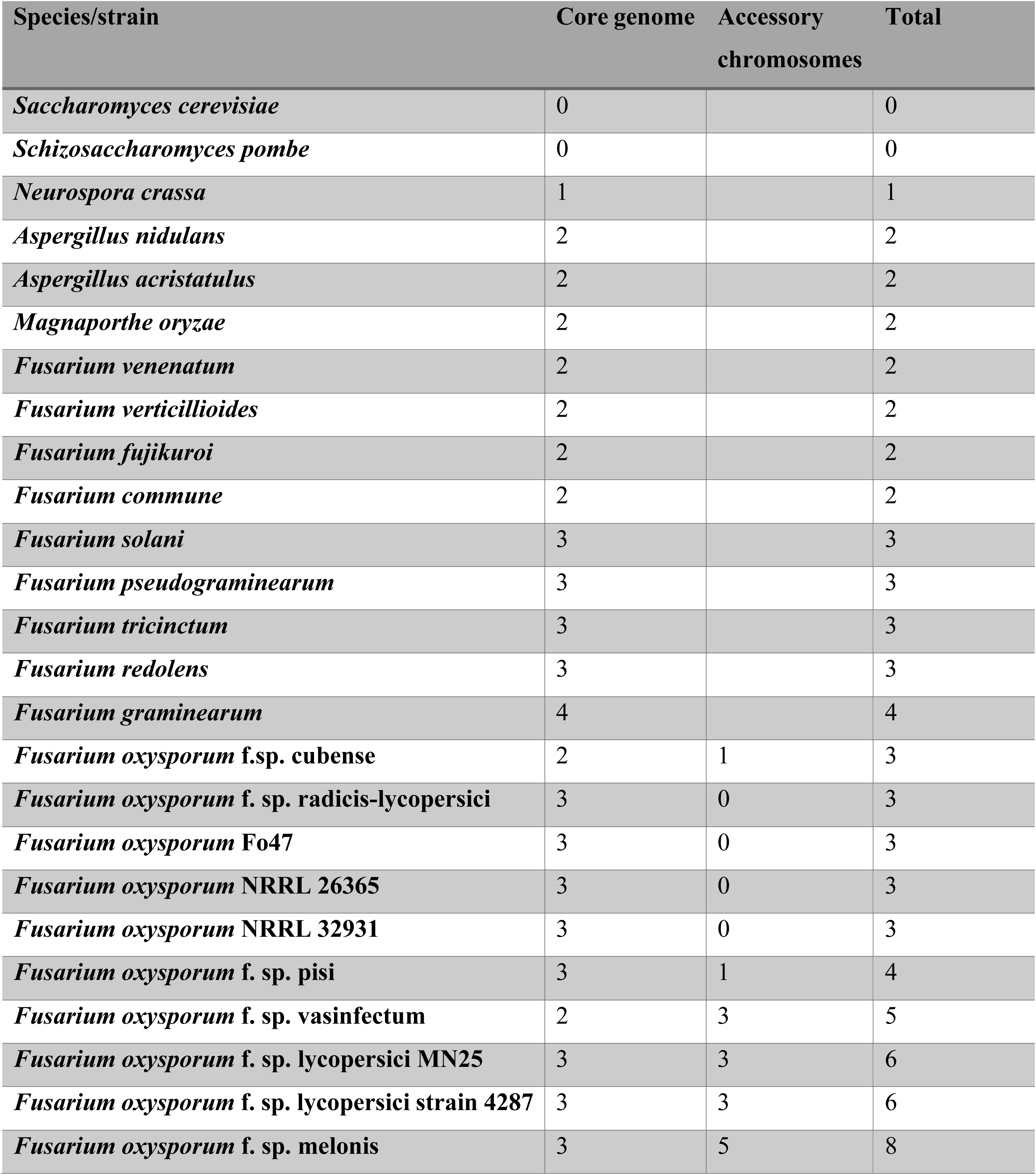
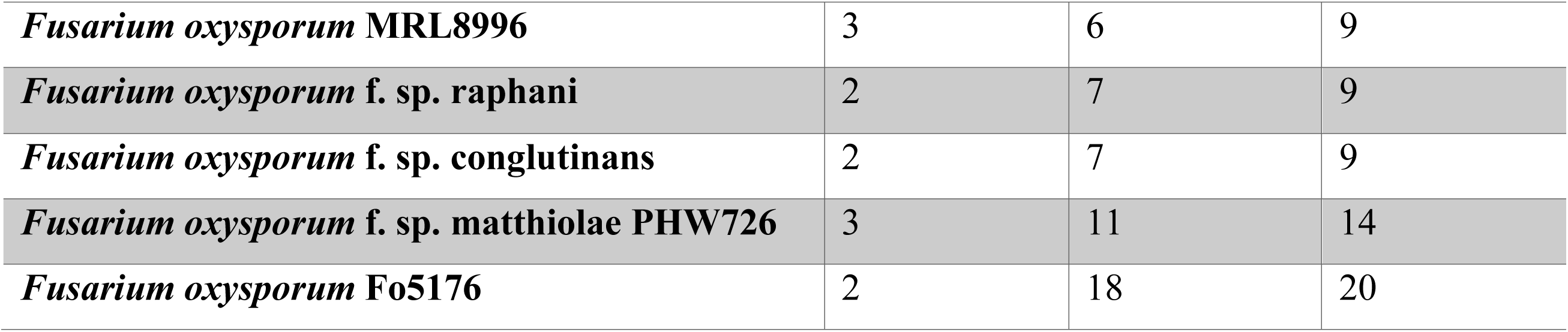
PARP copy number in 30 Ascomycetes.

Genome compartmentalization is a signature of a *F. oxysporum* genome. Core chromosomes (CCs) are conserved, vertically transmitted from parent to offspring, and involved in essential housekeeping functions, whereas accessory chromosomes (ACs) are thought to be horizontally transmitted, and are associated with specialized functions. Among FOSC strains, only up to three PARPs localized to CCs, while the remainder reside on ACs (Figure 3B). The three core genome encoding PARPs are a highly conserved PARP1 ortholog (>96% identity among the strains), a PARP-Ubc protein (>98% identity), and in some strains, a third short PARP catalytic domain-containing variant (“PARP-C”). In contrast, accessory PARPs vary widely in size and domain architecture; while all encode a variant of the catalytic domain, many lack the canonical WGR and regulatory domains. Notably, four strains (MRL8996, Fo5176, NRRL 26406, FoxPHW726) encode a complete PARP1 homolog on an accessory chromosome, in addition to the core copy.

### *Fusarium oxysporum* Accessory PARPs Reveal Convergent Evolution of Human-Like PARP Catalytic Motifs

To explore the functional diversity of accessory PARPs in the FOSC, we compared four strains with varying PARP repertoires. Fo47, a root-colonizing endophyte, represents the minimal configuration, encoding only three core PARPs and none on its ACs (Figure 3B). Fol4287, a tomato pathogen, and MRL8996, a human keratitis-infecting pathogen, each encode three core and three or six accessory PARPs, respectively (Figure 3B). Fo5176, an *A. thaliana* pathogen, harbors the largest known repertoire: two core and 18 accessory PARP proteins (Figure 3B).

The catalytic activity of PARPs depends on a conserved triad that is essential for NAD⁺ binding and chain elongation (Alemasova & Lavrik, 2019). Typically, the conserved DTX-family triad is H-Y-E in human poly(ADP-ribose) polymerases (hPARP1–5b) while Mono(ADP-ribosyl) transferases harbor alternative motifs (H-Y-I for hPARP7, H-Y-L for hPARP14, and H-Y-Y for hPARP16) (Vyas et al., 2014). Using human PARPs as references (Table S5), we identified the conserved H-Y-E triad in all core PARP1 and PARP-C homologs (Table 2). Notably, only 12 of the 38 accessory PARPs across the four FO strains contained catalytic residues (Table 2). All core PARP1 and PARP-C homologs retained the canonical H-Y-E motif. Three accessory PARPs (MRL8996_496866, Fo5176_269526, and Fo5176_525060) also contained intact H-Y-E triads and we identified six H-Y-I, two H-Y-L, and nine H-Y-Y variants in accessory PARPs. Interestingly, many FO PARPs co-harbored multiple motif variants (Table 2). For instance, all PARP1 and PARP-C homologs encode the H-Y-Y motif, in addition to the canonical H-Y-E motif (Table 2), while both motifs share the same H and Y residues. In fact, all cases of H-Y-Y variants co-exist with another catalytic triad, suggesting potential dual regulatory or structural roles. While detectable H-Y-E motif were present in nearly all FO PARPs, several showed partial loss of conserved residues (Table S6), implying potential loss of NAD⁺ binding or enzymatic function.

**Table 2.** Identified catalytic motifs in FO PARPs.

| FO strain | PARP protein | Identified catalytic motif<sup>[hPARP reference]</sup> |
| --- | --- | --- |
| FO47 | 216251 PARP1 | HYE<sup>[PARP1-5b]</sup>, HYY<sup>[PARP16]</sup> |
| FO47 | 274843 CC | HYE<sup>[PARP1-5b]</sup>, HYY<sup>[PARP16]</sup> |
| Fol4287 | 25845 PARP1 | HYE<sup>[PARP1-5b]</sup>, HYY<sup>[PARP16]</sup> |
| Fol4287 | 6509 CC | HYE<sup>[PARP1-5b]</sup>, HYY<sup>[PARP16]</sup> |
| Fol4287 | 9548 AC | HYI<sup>[PARP7]</sup>, HYY<sup>[PARP16]</sup> |
| Fol4287 | 13741 AC | HYI<sup>[PARP7]</sup>, HYY<sup>[PARP16]</sup> |
| MRL8996 | 468877 PARP1 | HYE<sup>[PARP1-5b]</sup>, HYY<sup>[PARP16]</sup> |
| MRL8996 | 516380 CC | HYE<sup>[PARP1-5b]</sup>, HYY<sup>[PARP16]</sup> |
| MRL8996 | 342866 AC | HYI<sup>[PARP7]</sup>, HYY<sup>[PARP16]</sup> |
| MRL8996 | 496866 AC | HYE<sup>[PARP1-5b]</sup> |
| Fo5176 | 329462 CC PARP1 | HYE<sup>[PARP1-5b]</sup>, HYY<sup>[PARP16]</sup> |
| Fo5176 | 432568 AC | HYI<sup>[PARP6]</sup> |
| Fo5176 | 525699 AC | HYE<sup>[PARP1-5b]</sup>, HYY<sup>[PARP16]</sup> |
| Fo5176 | 512474 AC | HYL<sup>[PARP14]</sup>, HYY<sup>[PARP16]</sup> |
| Fo5176 | 242882 AC | HYI<sup>[PARP7]</sup>, HYY<sup>[PARP16]</sup> |
| Fo5176 | 315425 AC | HYL<sup>[PARP14]</sup>, HYY<sup>[PARP16]</sup> |
| Fo5176 | 269526 AC | HYE<sup>[PARP1-5b]</sup>, HYY<sup>[PARP16]</sup> |
| Fo5176 | 525060 AC | HYE<sup>[PARP1-5b]</sup>, HYY<sup>[PARP16]</sup> |
| Fo5176 | 442380 AC | HYI<sup>[PARP6]</sup> |

We next screened for CTX-family motifs, using ectoART proteins as reference, and found no complete CTX motif (R-S-E) in the fungal PARPs. Interestingly, a potential S-S-S ectoART5 motif was identified in one of the accessory PARP variants (data not shown). However, although showing high proximity to the original positions, the residues in FO do not fully align with the reference and probably do not support the original enzymatic function. Taken together, these findings suggest that FO accessory PARPs have diverged from a DTX-family PARP1. However, while some retain recognizable catalytic motifs, others may be enzymatically inactive or functionally repurposed.

The identification of noncanonical human PARP catalytic motifs, some of which coexisting in the same accessory PARP protein, may indicate specific enzymatic adaptations that emerged independently across kingdoms of life. These findings allow us to exploit extensive knowledge of human PARPs to better understand fungal PARylation mechanisms and identify fungal-specific catalytic architectures with potential unique functions. To investigate the evolutionary origins of PARP catalytic motif diversity, we analyzed six fungal species representing major phyla: *Thelohania contejeani* (Microsporidia), *Synchytrium endobioticum* (Chytridiomycota), *Smittium culicis* (Zoopagomycota), *Gigaspora rosea* (Mucoromycota), *Gymnopus luxurians* (Basidiomycota), and *F. oxysporum* Fo5176 (Ascomycota). Using an rBBH and NCBI DELTA-BLAST pipeline, we identified all detectable PARP homologs in each species and aligned them against all human PARPs (Table S6). These sequences were also aligned to each other to generate a ML phylogenetic tree (Figure 4A). The resulting tree revealed two distinct clades. Clade A includes the Fo5176 PARP1 ortholog, all its accessory PARPs, two pairs of PARP homologs from Chytridiomycota and Mucoromycota, and one Basidiomycete PARP (Figure 4A). This confirms that FO accessory copies are PARP1 homologs. In contrast, Clade B includes Fo5176 PARP-Ubc, the Microsporidian PARP (harboring only H-Y-I motif), all Zoopagomycota PARPs (harboring both H-Y-E and H-Y-I motifs), and various Chytridiomycota, Mucoromycota, and Basidiomycota variants (Figure 4A). Interestingly, the H-Y-L motif was detected in Clade A, while absent from Clade B (Figure 4A).

**Figure 4.**
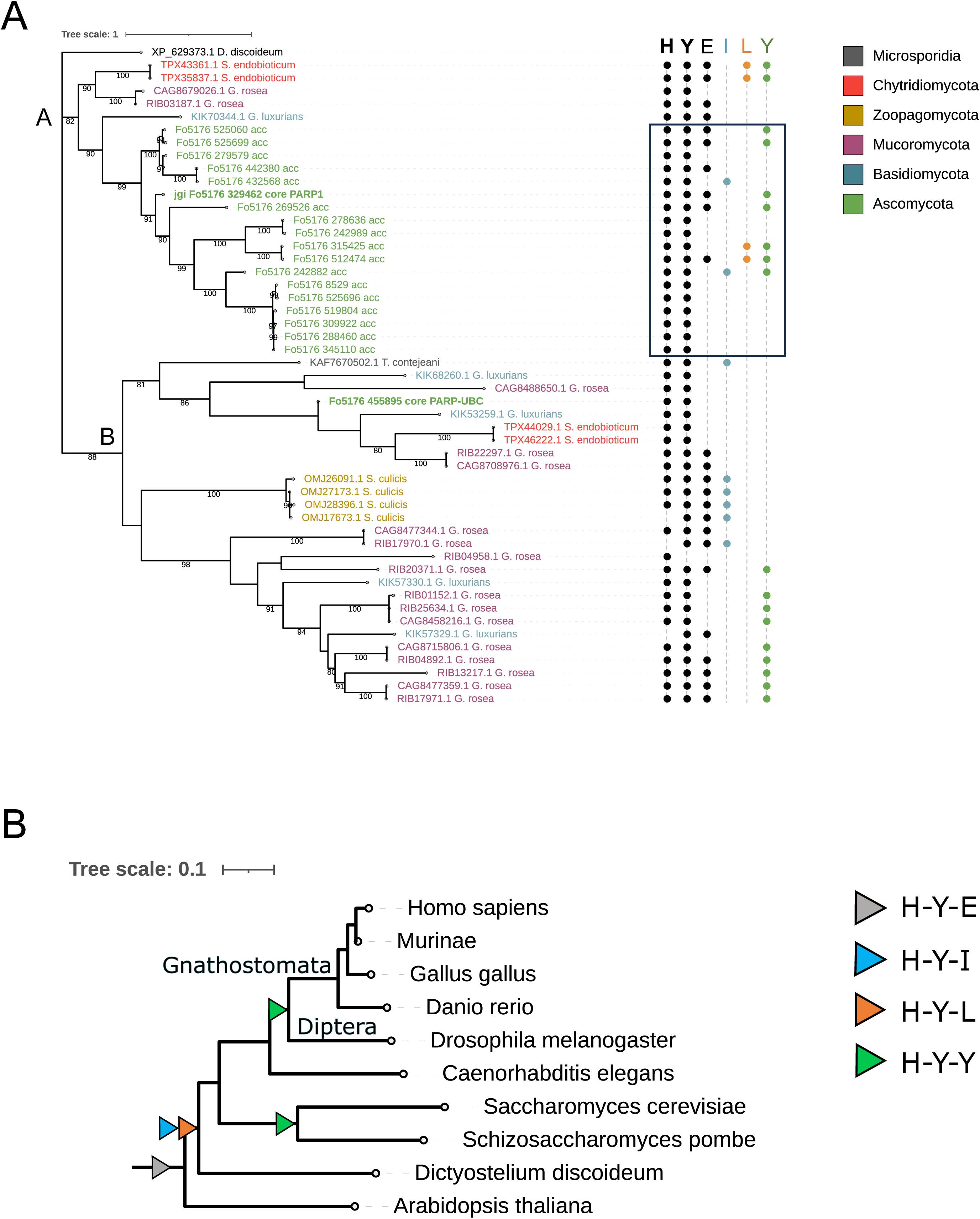
Evolutionary trajectories of the PARP catalytic motif variants in fungi A) Maximum likelihood tree of PARP protein repertoire of six fungal organisms, each represents a phylum within the fungi kingdom; *Thelohania contejeani* > Microsporidia, *Synchytrium endobioticum* > Chytridiomycota, *Smittium culicis* > Zoopagomycota, *Gigaspora rosea* > Mucoromycota, *Gymnopus luxurians* > Basidiomycota, *Fusarium oxysporum* Fo5176 > Ascomycota. Proteins were identified using reciprocal best BLAST hit and DELTA-BLAST. *Dictyostelium discoideum* served as an outgroup. Tree leaves are annotated with presence/absence of PARP catalytic triad variants (H-Y-E, H-Y-I, H-Y-L, H-Y-Y). Bootstrap values > 0.8 inferred from 1000 replicates are shown below the branches. All Fo5176 accessory PARPs are enclosed in a black square. B) The Tree of Life, generated by ITOL (Letunic & Bork, 2021), was pruned, visualized and annotated with emergence events of PARP catalytic triad variants. Tree includes Eukaryotic representatives from Plants, Amoebae, Fungi and Metazoa sharing last common ancestor at each node. The catalytic triad was identified by analyzing sets of multiple sequence alignment using all identified human PARPs as reference. Triangles at specific nodes are color- coded and indicate the presence of specific catalytic triad in the last common ancestor of the respective diverged branches.

To characterize the evolutionary variations in the PARP catalytic residues, we reconstructed the ancestral state of the identified catalytic motifs across eight representative eukaryotes from major lineages of Plants, Amoebae, Fungi and Metazoa (Figure 4B, Table S6). The canonical H-Y-E (DTX-family) motif, originally described in bacterial toxins, was detected in all PARP1 orthologs, including in *Dictyostelium discoideum*, which shares a common ancestor with both fungi and animals. Interestingly, *D. discoideum* also encodes additional PARPs with H- Y-I and H-Y-L motifs, corresponding to human PARP6 and PARP15, respectively. Along the metazoan lineage, the H-Y-Y motif (found in human PARP16) was identified in *Drosophila melanogaster*, *Danio rerio*, *Gallus gallus*, and humans, but not in *C. elegans*, suggesting its origin in the last common ancestor of Diptera and Gnathostomata (Figure 4B).

Taken together, these findings support a model of convergent evolution of a catalytic motif in fungal and animal PARPs. The independent emergence of the H-Y-Y motif across kingdoms suggests selective pressure for specific catalytic capabilities and potential integration of distinct ADP-ribosylation functions within fungal lineages.

### PARP Gene Copy Number Correlates with DNA Damage Tolerance and PARylation in *F. oxysporum*

To assess potential functional roles of the expanded PARP family in the four FOSC genomes, we constructed a phylogeny using the PARP catalytic domain (Figure 5A), integrated catalytic motif resolution (Figure 5B), and predicted the subcellular localization using DeepLoc 2.0 (Figure 5C, Table S7). Phylogenetically, all FO PARP repertoire clustered into two major clades. Clade A groups all PARP-Ubc genes, lacking a detectable catalytic motif (Figure 5A-B), yet all carry a nuclear localization signal (NLS) and are predicted to be exclusively nuclear (Figure 5C). Clade B includes all remaining PARPs, including PARP1, PARP-C, and accessory copies (Figure 5A). This clade is further subdivided into smaller sub-clades; however, bootstrap support is low and does not allow confident resolution of their evolutionary relationships. All PARP1 homologs are predicted to be nuclear (Figure 5C) and uniformly retained the canonical H-Y-E catalytic triad, whereas accessory PARPs frequently harbor catalytic variants and NLS, with most predicted to localize to the nucleus or both nucleus and cytoplasm (Figure 5B-C, Table S7). One exception, Fo5176_9589, carried both an NLS and a nuclear export signal (NES), consistent with mixed subcellular localization (Figure 5C, Table S7). Collectively, these results indicate that all FO accessory PARPs are more closely related to PARP1 orthologs than to PARP-Ubc, suggesting diversification from a canonical PARP1 ancestor (Figure 5A).

**Figure 5.**
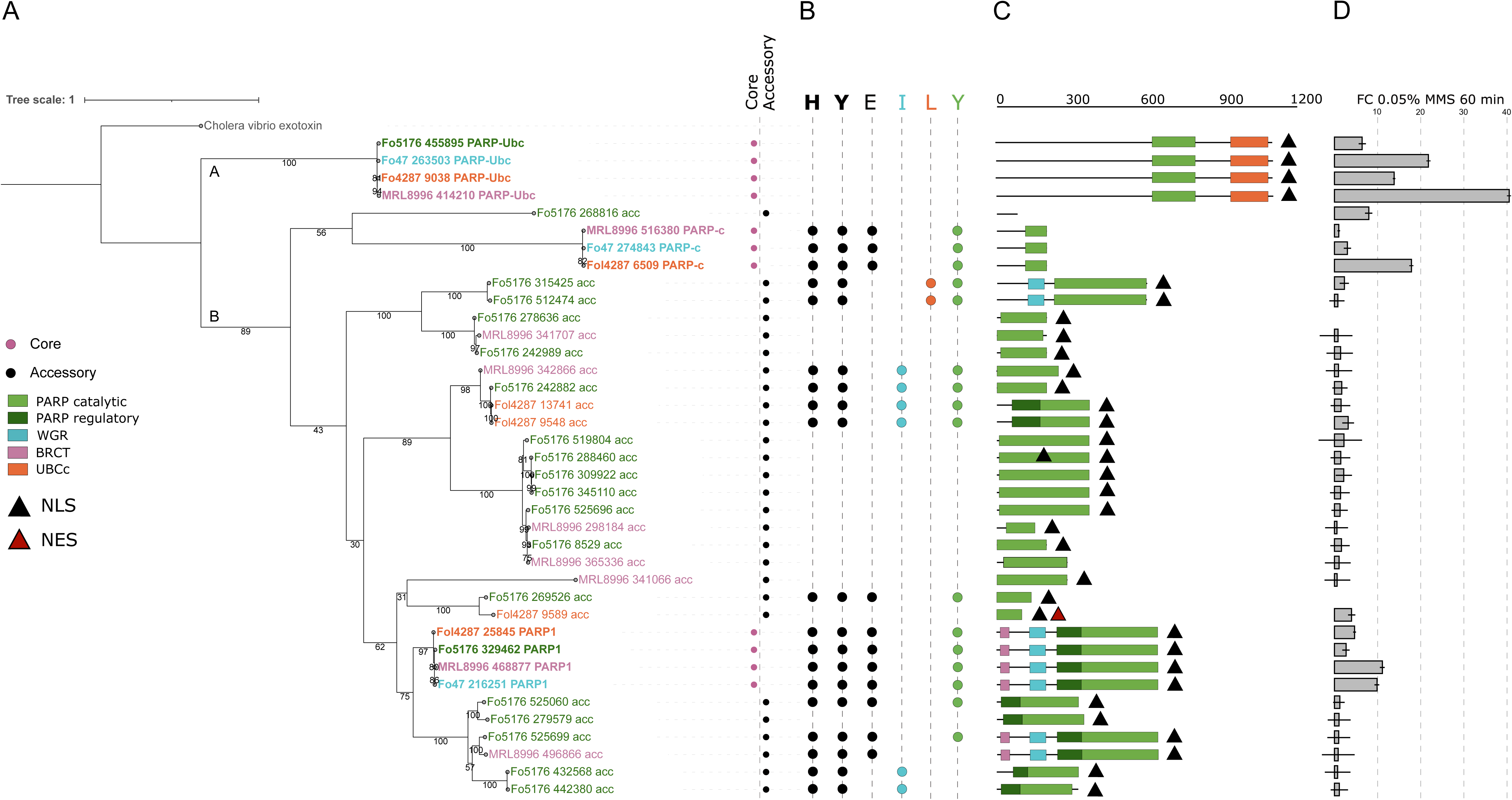
Characterization of PARP protein repertoire in the *F. oxysporum* species complex A) Maximum likelihood tree of PARP protein repertoire of four FO strains, Fo47 (turquoise), Fol4287 (orange), MRL8996 (pink), and Fo5176 (green). Bootstrap values inferred from 1000 replicates are shown below the branches. B) Identified catalytic motif of each PARP protein, Black circle indicates each residue of the identified catalytic motif. C) Conserved domain structure as inferred by NCBI CD-Search and InterProScan (Jones et al., 2014), nuclear localization signal (NLS) and nuclear export signal (NES) are indicated by black and scarlet triangle, respectively. D) qRT-PCR gene expression under DNA damage conditions (0.05% MMS, 60 minutes). Bar chart shows fold change values in treated cells compared to untreated control.

Since not all PARP proteins were predicted to harbor a catalytic motif or localization signal, we next assessed whether they are transcriptionally active using qRT-PCR under a treatment of methyl methanesulfonate (MMS), a DNA alkylating agent that activates the BER pathway (Figure 5D). *Parp1* expression increased in all strains to 9.8-, 4.6-, 11.1-, and 2.7-fold in Fo47, Fol4287, MRL8996, and Fo5176, respectively. *Parp-Ubc* was even more strongly induced to 21.7, 13.8, 40.5, and 6.4-fold in Fo47, Fol4287, MRL8996 and Fo5176, respectively. *Parp-C* was mildly induced in MRL8996 (3.0-fold), strongly in Fol4287 (17.8-fold), and showed no response in Fo47. Several accessory PARPs also showed strain-specific induction: Fol4287_9589 and 9548 (3.9- and 3.1-fold, respectively), and four Fo5176 genes (268816, 315425, 519804, 309922) showed 2.1-7.9-fold increases. No accessory genes in MRL8996 responded to MMS. These findings indicate that at least some accessory *Parp*s are transcriptionally active under DNA damage conditions.

Interestingly, we observed a negative correlation between the MMS-induced *Parp1* and *Parp-Ubc* expression and the number of transcriptionally active accessory *Parp*s across strains (r = -0.96 and -0.84, respectively, Figure S1). This inverse relationship was most evident in Fo5176, which harbors the highest number of active accessory PARPs and showed the lowest induction of both *Parp1* and *Parp-Ubc*, and, conversely, in MRL8996, which lacks any active accessory PARPs and exhibited the strongest induction of both genes (Figure 5D, S1). This finding suggests that active accessory *Parp*s may be part of a regulatory network that influences the expression of core *Parp* genes, potentially through feedback or compensatory mechanisms in response to DNA damage.

Because multiple accessory PARPs were transcriptionally active and displayed strain-specific expression patterns during MMS treatment, we next asked whether expansion of the PARP repertoire is associated with altered DNA damage tolerance. We, therefore, measured survival rates under genotoxic agents using our comparative system of four FO strains: Fo47, Fol4287, MRL8996, and Fo5176. We selected four DNA damage agents, including hydrogen peroxide and MMS, both of which induce single-strand breaks that are repaired via the BER pathway, 4- nitroquinoline oxide (4-NQO) that mimics UV lesions that are exclusively repaired by the NER pathway, and phleomycin (PLM) that induces DSBs. As shown in Figure 6A and 6B, strains with the highest copy numbers (MRL8996, Fo5176) displayed significantly reduced sensitivity to hydrogen peroxide and MMS (p < 10^−4^), while low-copy strains (Fo47, Fol4287) were more sensitive (Figure 6B-C). Differences under 4-NQO and PLM were significant but less pronounced (p < 10^⁻3^) (Figure 6B-C). This suggests that strains with higher PARP copy number are more resistant to single-strand breaks.

**Figure 6.**
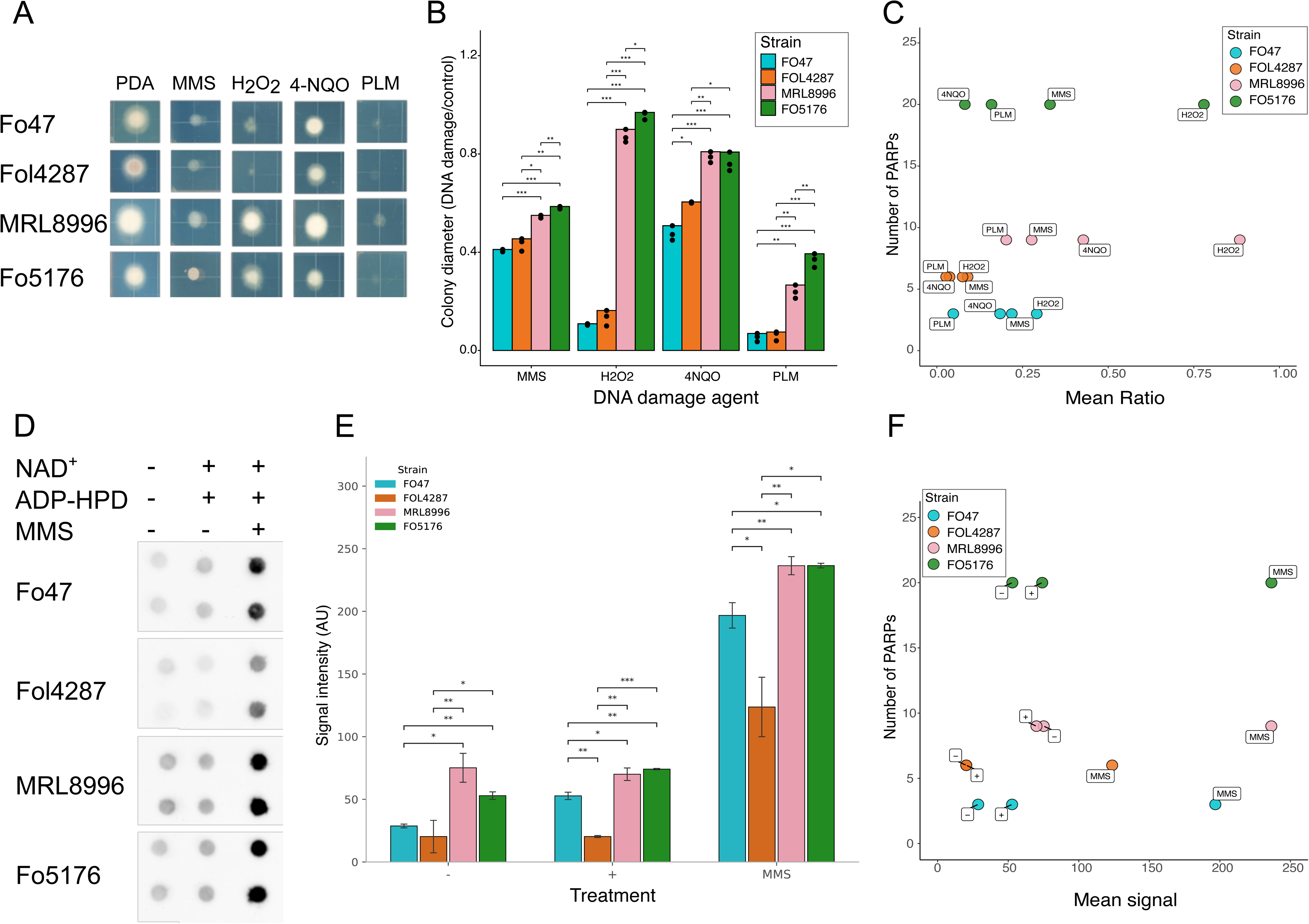
PARP copy number positively correlates with DNA damage tolerance and DNA damage- induced PARylation in *F. oxysporum* A) DNA damage survival assay of four FO strains: Fo47, Fol4287, MRL8996, and Fo5176. Strains were grown for 48 hours on DNA damage agent-containing PDA plates (0.01% MMS, 0.8 mM/mL H2O2, 0.45 ug/mL 4-NQO, and 0.4 ug/mL phleomycin). B-C) For each strain, colony diameter of three biological replicates of each condition were quantified using ImageJ. Significant differences of the ratio between mean values under DNA damage and control conditions were analyzed using two-way student t-test, * indicates p-value < 0.01, ** indicates p-value < 10^−4^. D) Quantification of cellular PARylation in Fo47, Fol4287, MRL8996, and Fo5176. Strains were incubated in normal conditions (PDB), PARylation-promoting conditions (PDB, 1 mM NAD^+^ and 2 μM of ADP-HPD) and DNA damage conditions (PDB, 1 mM NAD^+^, 2 μM of ADP-HPD and 0.05% MMS) for 20 minutes and were subjected to total protein extraction. Total cellular PARylation of proteins was determined by Immunodot blot assay using primary antibody against PAR (see materials and methods). E-F) Signal intensity of the Immunodot blot was quantified using ImageJ and mean values of raw intensity units for three biological replicates are shown. In both panels, “-”, “+” and “MMS” represent normal conditions, PARylation-promoting conditions, and DNA damage conditions, respectively.

Given these results, we asked whether PARP copy number correlates with basal PARylation activity. Using an immunodot blot assay with PAR-specific antibodies, we quantified PARylated protein levels in untreated and MMS-treated conidia, with or without PARP activation conditions: NAD⁺ + PARG inhibitor (PARGi). Under basal conditions (-), high-copy strains, MRL8996 and FO5176, displayed significantly elevated PAR signal compared to low-copy strains, FO47 and FOL4287 (FO47 vs. MRL8996: p=0.019; FO47 vs. FO5176: p=0.001; FOL4287 vs. MRL8996: p=0.006; FOL4287 vs. FO5176: p=0.043; Welch t-test), while the two high-copy strains did not differ significantly from each other (p=0.071), nor did the two low-copy strains (p=0.376). Following NAD⁺ + PARGi treatment (+), this separation was maintained and strengthened, with all low-copy vs high-copy comparisons reaching significance (p≤0.012), and FOL4287 vs FO5176 reaching p<0.001. The two high-copy strains did not differ significantly (p=0.291). Under MMS-induced DNA damage, high-copy strains showed significantly elevated PAR signal relative to both low-copy strains (p≤0.021), while MRL8996 and FO5176 remained statistically indistinguishable (p=0.989).

Taken together, our findings suggest that expanded PARP gene families contribute to enhanced tolerance to BER-associated DNA damage in *F. oxysporum*, potentially by sustaining higher levels of cellular PARylation. Accessory PARPs may contribute directly to this phenotype or indirectly through regulation of core PARP activity or other components of the DNA damage response. Their localization on accessory chromosomes further raises the possibility that these functions are particularly relevant under host-associated conditions.

## DISCUSSION

The PARP protein family is well characterized in animals due to its critical cellular functions and therapeutic potential (Kaur et al., 2022). Here, we present a comprehensive comparative genomic and evolutionary analysis of PARP proteins across the fungal kingdom. Our data show that the conservation of fungal PARP1 (hPARP1 homolog) and fungal PARP_Ubc (hPARP6 homolog) in fungi is highly dynamic. The fungal PARP1 gene was proposed to present in the last common ancestor of fungi. The ancestral-state reconstruction suggested that the fungal PARP6 gene (PARP-Ubc) was derived from multiple independent gain events based on its absence in the basal lineages, Microsporidia and Chytridiomycota. As a result, early diverging fungi rarely encode both PARP1 and PARP-Ubc, while their co-occurrence is common in filamentous Ascomycetes (specifically Pezizomycotina), which generally exhibit larger PARP repertoires, similar to other lineages across the tree of life (Perina et al., 2014).

A few facts, including the absence in the basal lineages, the fusion of a PARP6 catalytic domain and a UBCc domain only present in fungi (Citarelli et al., 2010), and the chytrid PARP6 homologs lack the UBCc domain suggests that the modern PARP-Ubc fusion emerged later in fungal evolution, along the Mucoromycota lineage. However, this inference should be interpreted cautiously given the extensive genome reduction and gene loss characteristic of Microsporidia. Moreover, homologs form a single phylogenetic clade, arguing against repeated independent emergence of the same PARP-UBCc fusion. Together with previous evidence that PARP6-like homologs may have been present in the last common ancestor of eukaryotes, these observations support an ancient origin of the PARP-Ubc lineage followed by differential retention and lineage-specific loss across fungi. In contrast to previous reports (Citarelli et al., 2010), we identified fungal species encoding PARP-Ubc in the absence of PARP1, primarily among Basidiomycetes and Ascomycetes. While previous work reported the absence of PARP proteins in most yeast lineages (Citarelli et al., 2010), our analysis suggests that losses have occurred independently across diverse fungal habitats and lifestyles. Conversely, PARP1 is retained in certain yeast clades. Microbotryomycetes in the subdivision Pucciniomycotina of the Basidiomycota is an example of a class containing many species in the form of yeast (Schoutteten et al., 2023), yet retaining a PARP enzyme due to an independent gain event. The loss of PARP proteins in Taphrinomycotina and Saccharomycotina have been previously reported, as has the presence of a PARP1 ortholog in *Yarrowia lipolytica* (Citarelli et al., 2010; Perina et al., 2014). Our analysis confirms these observations and reveals additional complexity, suggesting genus- or order-level lineage-specific trajectories. *Y. lipolytica* is a dimorph oleaginous yeast with specific industrial and financial interests due to its ability to uptake unusual carbon sources and consequently produce special lipids (Nicaud, 2012). The question why this specific organism retained PAR activity, while all the rest included in the same sub-division did not, is intriguing and may have several important implications. Investigating PARP activity in such species may shed light on PAR-related mechanism in uncommon PARP-encoding organisms as well as elucidate the frequent loss of PARP proteins in yeast lifestyle.

The evolution of fungal PARP1 domain architecture reflects both functional adaptations and lineage-specific divergence. Microsporidia have undergone an extensive genomic reduction, resulting in a massive loss of protein families and typically shorter proteins than their fungal orthologs (Wadi & Reinke, 2020). Indeed, the microsporidian PARP1 ortholog is short and lacks the canonical H-Y-E motif, harboring only a H-Y-I variant implying mono-ADP-ribosylation activity. In contrast, the chytrid PARP1 contains both PARP catalytic domain and WGR domain, likely reflecting the ancestral fungal domain architecture. This architecture suggests potential ADP-ribose transfer and DNA-binding functions, but not necessarily direct involvement in DNA repair activity. The latter appears to have occurred in Chytridiomycota through the acquisition of a BRCT domain, involved in DNA repair and PARP1 autoregulation (Leung & Glover, 2011). Several Mucoromycota species encode PARP variants with Tankyrase-like architectures. However, as previously reported in similar nonanimal PARP proteins (Citarelli et al., 2010; Perina et al., 2014), these protein variants contain a WGR rather than SAM domain and are classified as PARP1 homologs. These proteins also harbor a fungal-specific CHROMO domain, potentially linking between PAR-dependent telomere maintenance (Perina et al., 2014; Sowa et al., 2022) and heterochromatin remodeling in fungi. The emergence of telomere capping maintenance in the last common ancestor of Zoopagomycota and Mucoromycota (Wu et al., 2022) may promote such a function. Ascomycota exhibit the greatest diversity of domain architectures fused to PARP1, suggesting broader functional integration (Teichmann et al., 2001) and increased importance of PARP-related pathways in this group. Notably, we did not detect canonical PAR reader domains such as the Macrodomain in fungal PARPs, although such domains exist in fungi. This suggests that “writer-reader” coupling in PAR metabolism may have emerged later, or that alternative fungal-specific domain combinations confer analogous functions, a hypothesis requiring further investigation. The emergence of human-like catalytic motifs in fungal PARPs, some coexisting within the same protein, points to convergent evolution of the PARP catalytic activity (Gherardini et al., 2007). Given the contrasting evolutionary trajectories of animals and fungi (Ocaña-Pallarès et al., 2022), shared enzymatic motifs may suggest overlapping functional constraints. Unlike animals, fungal PARPs frequently combine multiple catalytic variants within a single protein, indicative of broader substrate promiscuity (Tawfik, 2010). Moreover, although Microsporidia and Zoopagomycota lack PARP6, their PARP1 orthologs often harbor its H-Y-I motif, either alone or in addition to H-Y-E. The catalytic domain sequence of these proteins also phylogenetically cluster closer to PARP-Ubc, despite structurally resembling PARP1. This suggests that at least one additional form of PARP, which does not fit into the distinct classification of a PARP1/PARP6 homolog, is present in fungi and may combine PARP1 structural features with catalytic motif typically associated with mono(ADP-ribosyl) transferases.

The PARP gene family have been considerably expanded during the evolution of Metazoa. The human PARP repertoire consists of 17 family members that were shaped to carry out and support a broad range of cellular activities (Vyas et al., 2013). Plants generally show smaller PARP family, while high copy number of PARP1/2 have been implicated to enhance tree longevity (Aoyagi Blue et al., 2021). We revealed a unique expansion of the PARP family in *Fusarium oxysporum* with expanded genes exclusively located in the lineage specific chromosomes. ACs in FO tend to harbor pathogenicity-related genes (Ma et al., 2010; Yang et al., 2020). The observed AC-driven expansion of PARP family implies a potential link between PAR-related activity and FO-specific pathogenic features. Our catalytic domain-based phylogeny suggests that all expanded copies are structurally similar to the PARP1 ortholog and therefore may possess similar functions even though many encode motif variants (H-Y-I, H-Y-L, H-Y-Y) that likely preclude full PARylation activity. It does not, however, resolve the precise evolutionary mechanism responsible for their expansion or exclude contributions from horizontal transfer. Addressing these questions will require broader phylogenomic sampling and synteny analyses. Although our dataset is limited in FO strain diversity, we noted that strains with high PARP copy numbers are Brassicaceae infecting pathogens. Given that camalexin, a Brassicaceae-derived ROS inducer, activates BER (Smith et al., 2013), additional PARP copies may enhance BER-mediated repair. Supporting this, several FO accessory PARPs were transcriptionally induced under MMS treatment, and “high copy number” strains exhibited significantly higher survival rates under H2O2 and basal PARylation levels under MMS conditions. Enhanced H2O2 resistance in MRL8996 compared with Fol4287, as recently shown by Ayhan et al. (2025) further supports our findings. Interestingly, Fol4287 and MRL8996 encode an equal number of catalytically complete accessory *Parp*s (four) yet show differential PARylation and DNA damage resistance. This suggests that not all catalytically intact copies contribute equally, or that differential PAR turnover may play a role despite the presence of a single PARG homolog in FO (Araiza-Cervantes et al., 2018). In line with this, we observed a negative correlation between the number of active accessory *Parp*s and the MMS-induced expression of core *Parp* genes, raising the possibility that accessory *Parp*s may influence core PARP activity through regulatory crosstalk or redundant mechanisms.

Our study provides a detailed map of PARP conservation, domain evolution, and expansion across the fungal kingdom. Yet, this is only the ‘tip of the iceberg’ in understanding the roles of PARP proteins in fungal physiology and pathogenicity. Studying these questions in an evolutionary context highlights the importance of PAR metabolism in fungi and the potential it poses as a target for novel antifungals. The wealth of currently available and constantly accumulating knowledge regarding PAR-related mechanisms in animals, offers a unique opportunity for the development of fungal-specific PARP inhibitors. The data presented here provides a good starting point to further explore PARylation and PARP related mechanisms in fungi.

## MATERIALS AND METHODS

### Strains and growth conditions

Four FO wild-type strains were used: Fo47, Fol4287, MRL8996, and Fo5176, and are described in more detail in Table S3. The *Fusarium oxysporum* strains *F. oxysporum* f. sp. *lycopersici* (Fol4287), *Fusarium oxysporum* FO47 (Fo47), *Fusarium oxysporum* MRL8996 (MRL8996) and *Fusarium oxysporum* Fo5176 (Fo5176) were chosen for all experiments due to their differential PARP family gene copy numbers. Fo47, Fol4287, MRL8996 and Fo5176 cultures were grown in sterile KNO3 based liquid medium and incubated at 28oC with shaking at 250 rpm for five days. The cultures were filtered to obtain microconidia that were washed in autoclaved ddH2O. Spores were dissolved in 1 mL autoclaved water and concentrations were determined using a hemocytometer. In all cases, spores were kept no longer than 24 hours at 4oC until further use.

### Identification of fungal PARP orthologs

Translated genomic sequence files were downloaded in FASTA format from the NCBI on August 10, 2022 and were used to construct a local BLAST database. The analysis included 534 species from nine phyla (Ascomycota, Basidiomycota, Blastocladiomycota, Chytridiomycota, Cryptomycota, Microsporidia, Mucoromycota, Rozellomycota and Zoopagomycota) within the fungal kingdom (Table S1 includes the complete list of proteome data sets). Orthologous genes were determined based on a reciprocal best BLAST hit (rBBH) approach (Tatusov, 2000) using in-house PERL scripts (Milo et al., 2019). To identify PARP1 orthologs, two different queries with two different seeds were conducted: First, *A. nidulans* PrpA (XP_660733.1), a previously identified PARP1 ortholog (Semighini et al., 2006), was used, followed by a query using the Fol4287 putative PARP1 ortholog (XP_018243277.1). All queries were executed against a total of 534 proteome files of fungal species (Table S1). The following parameters were used as the BlastP cutoff: E value of 0.1, length ratio of 0.45, and identity fraction of 0.25. Results were manually inspected and analyzed, including cases where no rBBH was found with BlastP. For PARP1, additional inspection was applied on cases where any discrepancy between the queries utilizing different seeds was observed. The orthologs of PARP1 and PARP-Ubc genes in all species are presented in Table S2.

### Gain and Loss analysis

Gain loss mapping engine (GLOOME) was used to predict gain and loss events of PARP genes based on the phyletic pattern and the phylogenetic tree (Cohen et al., 2010; Cohen & Pupko, 2011). For the estimation of the expectation of the number of gain and loss events for each *Parp* gene over the evolutionary tree, the optimization level was set to ‘very high’, a fixed gain/loss model was used, and the distribution type was ‘Gamma’. Events with probability value ≥ 0.50 were extracted (Table S2). The sum of expectation values for gain and loss events of a gene is termed here the GLOOME score.

### Identification of PARP family of protein in the FOSC

Proteomes of 30 fungal species (Table S3), representing major Ascomycete classes (Saccharomycetes, Schizosaccharomycetes, Eurotiomycetes, and Sordariomycetes) with different lifestyles (yeast, filamentous fungi, saprophytic and pathogenic) were downloaded from JGI MycoCosm portal (https://mycocosm.jgi.doe.gov/mycocosm/home) (Grigoriev et al., 2014) (Table S3). To identify the PARP family, protein sequence of the Fol4287 PARP1 ortholog (XP_018243277.1) was downloaded and submitted to ENSEMBL BioMart (https://fungi.ensembl.org/biomart) using the ENSEMBL Fungi Genes 54 database and *Fusarium oxysporum* genes (FO2) dataset. A list of InterPro protein domains and/or families was retrieved and processed by a custom R script to search for all annotated proteins that harbor at least one PARP domain/family module, resulting in a list of identified protein families within each included species (Table S4).

### Genome partition

Genome partition was retrieved from previous reports for Fol4287 (Ma et al., 2010), FoII5 (YONG ZHANG, 2019), Fo5176 (Fokkens et al., 2020) and Fo47 (B. Wang et al., 2020). Fo47 shows clear genome partition with 11 core and one accessory chromosome and therefore served as the reference for this matter. The rest eleven *F. oxysporum* genomes were partitioned into core and accessory chromosomes using mummer/3.22 (Kurtz et al., 2004) by aligning scaffolds of the genome assemblies against the core chromosomes of the reference Fo47 with default parameters. The scaffolds aligned to the core chromosomes of Fo47 showing coverage larger than 5% were annotated as core scaffolds while all the rest were considered as accessory scaffolds.

Conserved domain analysis of fungal PARP1 and PARP-Ubc NCBI CD-Search and Batch CD-search (https://www.ncbi.nlm.nih.gov/Structure/cdd/cdd.shtml) were used to search the Conserved Domain Database (CDD v3.19 – 58235 PSSMs) using PARP protein sequences in FASTA format with the following parameters: E value threshold of 0.01 and “Composition based statistics adjustment” set to “yes”. In several cases, additional annotation was performed using InterProScan/5.38-76.0 (https://www.ebi.ac.uk/interpro/search/sequence) (Jones et al., 2014). All protein schematic models were drawn manually based on the CDD-search or InterProScan coordinates.

### Computational prediction of catalytic motifs in PARP homologs

Amino acid sequences FO PARPs were retrieved from the JGI MycoCosm portal (https://mycocosm.jgi.doe.gov/mycocosm/home) (Grigoriev et al., 2014). PARP homologs of Fungal (*Thelohania contejeani* > Microsporidia, *Synchytrium endobioticum* > Chytridiomycota, *Smittium culicis* > Zoopagomycota, *Gigaspora rosea* > Mucoromycota, *Gymnopus luxurians* > Basidiomycota) and other Eukaryotic (*Dictyostelium discoideum* AX4, *Caenorhabditis elegans*, *Drosophila melanogaster*, *Danio rerio*, *Gallus gallus*) representatives were identified using a combination of rBBH and DELTA-BLAST searches. Results were manually inspected, and duplicates were removed. Amino acid sequences of all identified PARP homologs were retrieved from NCBI Genbank (Benson et al., 2013). Amino acid sequences of *Homo sapiens* PARP proteins were located using accession numbers compiled by Hottiger et al. Catalytic domains of all organisms were identified using InterPro Search (Blum et al., 2021) and retrieved using a custom R script. Each set of PARP sequences per organism were repeatedly aligned against each human PARP reference using the Multiple Alignment using Fast Fourier Transform (MAFFT) program (version 7.313) (Katoh & Standley, 2013) with default parameters. The output format of ClustalW with character counts was visualized using MView (https://www.ebi.ac.uk/Tools/msa/mview/) with default parameters. Human catalytic motif residue locations were manually annotated in the multiple sequence alignment (MSA) results (Otto et al., 2005) and identified catalytic residues were documented in Table S6.

### Subcellular localization of FO PARP proteins

Subcellular localization for all PARP proteins included in our four-strains comparative system was determined using DeepLoc 2.0 (Thumuluri et al., 2022) with ‘High-quality’ model (Table S7).

### Construction and visualization of phylogenetic trees

For the fungal tree depicting the conservation of PARP1 and PARP-Ubc orthologs (Figure 1) and diversity of PARP1 architecture (Figure 2) across the fungal kingdom, a genome-scale tree was adopted from (Li et al., 2021). The tree was pruned using the ‘Ape’ R library (version 5.6-2) to a final version of 534 fungal species. To visualize a tree showing evolutionary trajectories of PARP catalytic motifs in Eukaryotes (Figure 4B), the same approach was used using the Tree Of Life by Interactive Tree of Life (ITOL, https://itol.embl.de) (Letunic & Bork, 2021) as a template. A total of 500 single-copy orthologs determined by OrthoFinder 2.5.4 (Emms & Kelly, 2019) were used to construct the Ascomycetes (Figure 3A) and FO strains (Figure 3B) trees. The FO tree was rooted by *F. verticillioides*. Trees showing fungal PARP family (Figure 4A) and FO PARP repertoire (Figure 5A) were constructed using amino acid sequences of the PARP catalytic domain. All trees were aligned using MAFFT (version 7.313) (Katoh & Standley, 2013) with default parameters and output was used to construct maximum likelihood (ML) trees by IQ-TREE version 1.6.3 (-bb 1000 -ntmax 4 -st AA -wbt -alrt 1000) (Nguyen et al., 2015). Bipartition values of the best ML tree were estimated. In some cases, multiple sequence alignments were processed using noisy 1.5.12 (Dress et al., 2008) to identify problematic columns before submitted to IQ-TREE. All phylogenetic trees were visualized and annotated in ITOL (https://itol.embl.de) (Letunic & Bork, 2021) using the ITOL Annotation Editor (https://itoleditor.letunic.com).

### Transcriptional response to DNA damage

Three biological replicates containing 10^8^ spores of FO each were incubated in 10mL of PDB media (19.2g of Potato Dextrose Broth, 800mL of water) in an orbital shaker at 200 rpm at 28℃ for 3 hours. Strains used (Fo47, Fol4287, MRL8996, and Fo5176,) are described in Table S3. Samples were removed from the shaker, and two groups of samples were created: a control group with no additional added reagents, and a treated group supplemented with 0.05% MMS. All samples were reinserted into the orbital shaker at the same rpm and temperature for 1 hour of additional incubation. Collected samples were centrifuged at 4000 x g at 4°C for 10 minutes, flash frozen with liquid nitrogen and stored at -80℃ overnight.

### RNA extraction

Conidia were disrupted in a temperature-controlled tissue homogenizer (Bullet Blender, Next Advance Inc, NY, U.S.A) for 90 seconds at maximum speed. Total RNA was extracted using the Qiagen RNeasy micro kit (California, U.S.A) and was treated on-column with RQ1 RNase-free Dnase (Promega; Wisconsin, U.S.A). Each RNA sample was quantified in a Nanodrop ND- 1000 UV-Vis Spectrophotometer (Thermo Fisher Scientific; Massachusetts, U.S.A) using 1 µl RNA.

### cDNA synthesis and quantitative PCR expression analysis

cDNA was generated using Qiagen Omniscript RT kit (Qiagen; California, U.S.A) according to the manufacturer’s instructions using 1 μg extracted RNA. Quantitative PCR was carried out using Fast SYBR® Green Master Mix (Thermo Fisher Scientific, Massachusetts, U.S.A). Each well contained 2 ng cDNA. The final reaction volume in each well was 10 μl in 0.1 ml volume 96-well plates (USA Scientific; Florida, U.S.A). The reactions were carried out on an Eppendorf MasterCycler EpGradient realplex2 real-time PCR machine (Eppendorf; Connecticut, U.S.A). Raw data were downloaded and analyzed in MS Excel (Microsoft; Washington, U.S.A). All data were compared using the comparative ΔΔ*CT* method (Pfaffl, 2001). All primer pairs (Table S8) amplified their target with equal efficiency (data not shown).

### DNA damage sensitivity assays and data analysis

For DNA damage sensitivity assays, freshly harvested microconidia was grown on PDA (BD Difco; Maryland, U.S.A) plates supplemented with either of the following DNA damage agents: Hydrogen peroxide (H2O2) Fisher Scientific; Massachusetts, U.S.A), methyl methanesulfonate (MMS) (Sigma-Aldrich; Missouri, U.S.A), 4-nitroquinoline oxide (4-NQO) (Sigma-Aldrich; Missouri, U.S.A) and phleomycin (PLM) (Sigma-Aldrich; Missouri, USA) at the indicated final concentration; 0.8 mM H2O2, 0.01% MMS, 0.45 μg/mL 4-NQO, and 0.4 μg/mL PLM. 10×10 square plates containing 30 mL of PDA medium and the required damage agent were freshly made on the same day of assay and allowed to solidify in the dark in a biological hood for several hours. 10 μL of spore suspensions of 10^6^, 10^5^, and 10^4^ spores/mL concentrations were inoculated in three replicates on the same media plates for a final concentration of 10^4^, 10^3^, and 10^2^ spores-containing colonies. The plates were incubated in a complete dark at 28°C for 48 hours and scanned. Colony size was quantified using ImageJ (version 1.53k). To quantify the effect of DNA damage tolerance, the values of the 10^3^ treated replicates were divided by the values of the 10^3^ control replicates followed by a two-way student’s t-test between the strains in all possible combinations.

### Experimental design and conditions of protein PARylation induction

A total of 2.0×10^8^ spores of each strain were incubated in 10 mL of Potato Dextrose Broth (PDB) (BD Difco; Maryland, U.S.A) medium (19.2 g of Potato Dextrose Broth, 800 mL of water) in an orbital shaker at 200 rpm at 28°C for 3 hours. Samples were removed from the orbital shaker and were split into three groups: 1) a control group with no added supplements, 2) a group where 2 μM of ADP-HPD (Sigma-Aldrich; Missouri, U.S.A), and 1 mM of NAD^+^ (Sigma-Aldrich; Missouri, U.S.A) were added and 3) a group where 0.1% MMS, 2 μM of ADP-HPD, and 1 mM of NAD^+^ were added. All samples were reinserted into the orbital shaker for additional incubation of 20 minutes at the same rpm and temperature. Collected samples were centrifuged at 4000 x *g* at 4°C for 10 minutes, after which the supernatant was discarded, and samples were flash frozen in liquid nitrogen until further use. For protein extraction, samples were partially thawed at room temperature and immediately vortexed with 125 μL of lysis buffer [40 mM potassium phosphate buffer, pH 7.0, 5 mM Ethylenediaminetetraacetic acid (EDTA), 0.1% Triton X-100, 20% glycogen, 1 ug of Dichlorodiphenyltrichloroethane (DDT), and 10 uL of plant protease inhibitor (Sigma-Aldrich; Missouri, U.S.A) per mL]. Following lysis, the samples were sonicated using a Heat Systems W-385 Sonicator Ultrasonic Processor at 50% duty cycle for 1 second for 5 times; samples were then shaken at 50 rpm on an orbital shaker for 15 minutes and finally the samples were transferred to 1.5 mL Eppendorf tubes, and centrifuged at 13000 x *g* at room temperature for 10 minutes. The supernatant was transferred to a new 1.5 mL tube while the pellet was discarded. Total protein extracts were quantified using the Bradford method with Bio-Rad Protein Assay Dye Reagent Concentrate (Bio-Rad Laboratories; California, U.S.A) and Bovine Serum Album (Pierce Biotechnology; Illinois, U.S.A) as the standard for generating a standard curve; absorbance was measured at 595 nm wavelength. The total protein extracts were stored at -20°C.

### Immunodot blot assay

Protein samples were thawed on ice and then diluted to a concentration of 10 μg/mL. Two units of Bio-Rad Bio-Dot/Bio-Dot SF Filter Paper (Bio-Rad; California, U.S.A) were stacked with Amersham Hybond-N Membrane (GE Healthcare Limited; Buckingshire, England) and submerged in TBS (Tris-Buffered Saline, pH 7.4, Thermo Fisher; Ontario, Canada) for 10 minutes. The Dot-Blot Apparatus was assembled using the soaked membrane and filter papers, with the nylon membrane on top. A total volume of 225 μL of each sample was blotted onto the membrane and the same volume of PBST (Phosphate-Buffered Saline, pH 7.4, Thermo Fisher; Ontario, Canada with 0.1% Tween20, Fair Lawn; New Jersey, U.S.A) was added into empty wells. BSA (10 μg/mL) was used as a negative control. A light vacuum was applied until all samples had been vacuumed through the membrane, followed by an additional 225 μL of PBST added to all wells. The membrane was vacuumed and then dried using a Model 583 Gel Dryer (Bio-Rad; California, U.S.A) for 90 minutes at 80°C. The membrane was rehydrated in TBST (Tris-Buffered Saline, pH 7.5, with 0.1% Tween20) for 5 minutes and then incubated in 10% formaldehyde solution for 20 minutes at 37°C. The membrane was washed on an orbital shaker with TBS twice for two minutes. Next, the membrane was placed between two filter papers and dried on a gel dryer for an additional hour at 80°C. The membrane was rehydrated with PBST for 5 minutes. From this point, all washing and incubation steps were performed on an orbital shaker at 60 rpm. The membrane was incubated with a blocking solution (5% Difco Skim Milk, Sparks; Maryland, U.S.A, in PBST) for one hour. The membrane was washed with PBST three times for 20 minutes each, with PBST. Next, the membrane was incubated with the primary antibody solution 1:1000 Poly(ADP-ribose) monoclonal antibody (10H) (Enzo Biochem Inc.; New York, U.S.A) in blocking solution overnight at 4°C. The following morning, the membrane was washed three times for 20 minutes each, with PBST. The membrane was then incubated with a secondary antibody solution 1:1000 Goat Anti-Mouse IgG Antibody, (H+L) HRP conjugate (EMD Millipore; Missouri, U.S.A) in blocking solution for one hour. The membrane was washed four times for 30 minutes in PBST. Next, 2.5 mL of Enhanced chemiluminescence (ECL) (Life Technology Corporation; Ontario, Canada) reagent 1 and 2 were mixed in a 1:1 ratio, pipetted evenly onto the membrane and allowed to incubate while covered for 2 minutes. The membrane was gently dried imaged for chemiluminescence in a BioRad ChemiDoc Imaging System (Bio-Rad; California, U.S.A). Signal was quantified using ImageJ (version 1.53k) where the mean background signal was calculated and subtracted from each of the sample measurements. The mean signal value of technical replicates was calculated and used for data points in further analysis.

### Statistical Analyses and Data Visualization

Data manipulation and analysis was done using Python 3.12.2 in JupyterLab (version 4.4.10) (https://jupyter.org). Data was plotted and statistically analyzed using R base functions and the ‘ggplot2’ R library (version 3.4.0). The scripts were executed in R studio console version 2022.12.0 Build 353 (https://www.rstudio.com) using R language version 4.1.2 (https://www.r-project.org).

## REFERENCES

Alemasova, E. E., & Lavrik, O. I. (2019). Poly(ADP-ribosyl)ation by PARP1: reaction mechanism and regulatory proteins. Nucleic Acids Research, 47(8), 3811–3827. 10.1093/nar/gkz120

Aoyagi Blue, Y., Kusumi, J., & Satake, A. (2021). Copy number analyses of DNA repair genes reveal the role of poly(ADP-ribose) polymerase (PARP) in tree longevity. IScience, 24(7), 102779. 10.1016/j.isci.2021.102779

Araiza-Cervantes, C. A., Meza-Carmen, V., Martínez-Cadena, G., Roncero, M. I. G., Reyna-López, G. E., & Franco, B. (2018). Biochemical and genetic analysis of a unique poly(ADP-ribosyl) glycohydrolase (PARG) of the pathogenic fungus Fusarium oxysporum f. sp. lycopersici. Antonie van Leeuwenhoek, 111(2), 285–295. 10.1007/s10482-017-0951-2

Ayhan, D. H., Abbondante, S., Martínez-Soto, D., Wu, S., Rodriguez-Vargas, R., Milo, S., Rickelton, K., Sohrab, V., Kotera, S., Arie, T., Marshall, M. E., Rocha, M. C., Haridas, S., Grigoriev, I. V., Shlezinger, N., Pearlman, E., & Ma, L.-J. (2025). Host-specific adaptation in *Fusarium oxysporum* correlates with distinct accessory chromosome content in human and plant pathogenic strains. MBio, 16(8). 10.1128/mbio.00951-25

Baroncelli, R., Cobo-Díaz, J. F., Benocci, T., Peng, M., Battaglia, E., Haridas, S., Andreopoulos, W., LaButti, K., Pangilinan, J., Lipzen, A., Koriabine, M., Bauer, D., Le Floch, G., Mäkelä, M. R., Drula, E., Henrissat, B., Grigoriev, I. V, Crouch, J. A., de Vries, R. P., … Thon, M. R. (2024). Genome evolution and transcriptome plasticity is associated with adaptation to monocot and dicot plants in *Colletotrichum* fungi. GigaScience, 13. 10.1093/gigascience/giae036

Benson, E., Betson, F., Fuller, B. J., Harding, K., & Kofanova, O. (2013). Translating cryobiology principles into trans-disciplinary storage guidelines for biorepositories and biobanks: a concept paper. Cryo Letters, 34(3), 277–312.

Blum, M., Chang, H.-Y., Chuguransky, S., Grego, T., Kandasaamy, S., Mitchell, A., Nuka, G., Paysan-Lafosse, T., Qureshi, M., Raj, S., Richardson, L., Salazar, G. A., Williams, L., Bork, P., Bridge, A., Gough, J., Haft, D. H., Letunic, I., Marchler-Bauer, A., … Finn, R. D. (2021). The InterPro protein families and domains database: 20 years on. Nucleic Acids Research, 49(D1), D344–D354. 10.1093/nar/gkaa977

Britton, T. A., Wu, C., Chen, Y.-W., Franklin, D., Chen, Y., Camacho, M. I., Luong, T. T., Das, A., & Ton- That, H. (2024). The respiratory enzyme complex Rnf is vital for metabolic adaptation and virulence in *Fusobacterium nucleatum*. MBio, 15(1). 10.1128/mbio.01751-23

Cho-Park, P. F., & Steller, H. (2013). Proteasome Regulation by ADP-Ribosylation. Cell, 153(3), 614–627. 10.1016/j.cell.2013.03.040

Citarelli, M., Teotia, S., & Lamb, R. S. (2010). Evolutionary history of the poly(ADP-ribose) polymerase gene family in eukaryotes. BMC Evolutionary Biology, 10(1), 308. 10.1186/1471-2148-10-308

Cohen, O., Ashkenazy, H., Belinky, F., Huchon, D., & Pupko, T. (2010). GLOOME: gain loss mapping engine. Bioinformatics, 26(22), 2914–2915. 10.1093/bioinformatics/btq549

Cohen, O., & Pupko, T. (2011). Inference of Gain and Loss Events from Phyletic Patterns Using Stochastic Mapping and Maximum Parsimony—A Simulation Study. Genome Biology and Evolution, 3, 1265–1275. 10.1093/gbe/evr101

Deeksha, W., Abhishek, S., & Rajakumara, E. (2023). PAR recognition by PARP1 regulates DNA- dependent activities and independently stimulates catalytic activity of PARP1. The FEBS Journal, 290(21), 5098–5113. 10.1111/febs.16907

DeIulio, G. A., Guo, L., Zhang, Y., Goldberg, J. M., Kistler, H. C., & Ma, L.-J. (2018). Kinome Expansion in the Fusarium oxysporum Species Complex Driven by Accessory Chromosomes. MSphere, 3(3). 10.1128/mSphere.00231-18

Dress, A. W., Flamm, C., Fritzsch, G., Grünewald, S., Kruspe, M., Prohaska, S. J., & Stadler, P. F. (2008). Noisy: Identification of problematic columns in multiple sequence alignments. Algorithms for Molecular Biology, 3(1), 7. 10.1186/1748-7188-3-7

Emms, D. M., & Kelly, S. (2019). OrthoFinder: phylogenetic orthology inference for comparative genomics. Genome Biology, 20(1), 238. 10.1186/s13059-019-1832-y

Feijs, K. L. H., Forst, A. H., Verheugd, P., & Lüscher, B. (2013). Macrodomain-containing proteins: regulating new intracellular functions of mono(ADP-ribosyl)ation. Nature Reviews Molecular Cell Biology, 14(7), 443–451. 10.1038/nrm3601

Fokkens, L., Guo, L., Dora, S., Wang, B., Ye, K., Sánchez-Rodríguez, C., & Croll, D. (2020). A Chromosome-Scale Genome Assembly for the *Fusarium oxysporum* Strain Fo5176 To Establish a Model *Arabidopsis* -Fungal Pathosystem. G3 Genes|Genomes|Genetics, 10(10), 3549–3555. 10.1534/g3.120.401375

Gao, X., Gao, G., Zheng, W., Liu, H., Pan, W., Xia, X., Zhang, D., Lin, W., Wang, Z., & Feng, B. (2024). PARylation of 14-3-3 proteins controls the virulence of Magnaporthe oryzae. Nature Communications, 15(1), 8047. 10.1038/s41467-024-51955-w

Gherardini, P. F., Wass, M. N., Helmer-Citterich, M., & Sternberg, M. J. E. (2007). Convergent Evolution of Enzyme Active Sites Is not a Rare Phenomenon. Journal of Molecular Biology, 372(3), 817–845. 10.1016/j.jmb.2007.06.017

Gibson, B. A., & Kraus, W. L. (2012). New insights into the molecular and cellular functions of poly(ADP-ribose) and PARPs. Nature Reviews Molecular Cell Biology, 13(7), 411–424. 10.1038/nrm3376

Grigoriev, I. V., Nikitin, R., Haridas, S., Kuo, A., Ohm, R., Otillar, R., Riley, R., Salamov, A., Zhao, X., Korzeniewski, F., Smirnova, T., Nordberg, H., Dubchak, I., & Shabalov, I. (2014). MycoCosm portal: gearing up for 1000 fungal genomes. Nucleic Acids Research, 42(D1), D699–D704. 10.1093/nar/gkt1183

Guo, X., Carroll, J.-W. N., MacDonald, M. R., Goff, S. P., & Gao, G. (2004). The Zinc Finger Antiviral Protein Directly Binds to Specific Viral mRNAs through the CCCH Zinc Finger Motifs. Journal of Virology, 78(23), 12781–12787. 10.1128/JVI.78.23.12781-12787.2004

Hottiger, M. O., Hassa, P. O., Lüscher, B., Schüler, H., & Koch-Nolte, F. (2010). Toward a unified nomenclature for mammalian ADP-ribosyltransferases. Trends in Biochemical Sciences, 35(4), 208–219. 10.1016/j.tibs.2009.12.003

Hsiao, S. J., & Smith, S. (2008). Tankyrase function at telomeres, spindle poles, and beyond. Biochimie, 90(1), 83–92. 10.1016/j.biochi.2007.07.012

Jones, P., Binns, D., Chang, H.-Y., Fraser, M., Li, W., McAnulla, C., McWilliam, H., Maslen, J., Mitchell, A., Nuka, G., Pesseat, S., Quinn, A. F., Sangrador-Vegas, A., Scheremetjew, M., Yong, S.-Y., Lopez, R., & Hunter, S. (2014). InterProScan 5: genome-scale protein function classification. Bioinformatics, 30(9), 1236–1240. 10.1093/bioinformatics/btu031

Kalisch, T., Amé, J.-C., Dantzer, F., & Schreiber, V. (2012). New readers and interpretations of poly(ADP-ribosyl)ation. Trends in Biochemical Sciences, 37(9), 381–390. 10.1016/j.tibs.2012.06.001

Katoh, K., & Standley, D. M. (2013). MAFFT Multiple Sequence Alignment Software Version 7: Improvements in Performance and Usability. Molecular Biology and Evolution, 30(4), 772–780. 10.1093/molbev/mst010

Kaur, S. D., Chellappan, D. K., Aljabali, A. A., Tambuwala, M., Dua, K., & Kapoor, D. N. (2022). Recent advances in cancer therapy using PARP inhibitors. Medical Oncology, 39(12), 241. 10.1007/s12032-022-01840-7

Kelkar, Y. D., & Ochman, H. (2012). Causes and Consequences of Genome Expansion in Fungi. Genome Biology and Evolution, 4(1), 13–23. 10.1093/gbe/evr124

Kleine, H., Poreba, E., Lesniewicz, K., Hassa, P. O., Hottiger, M. O., Litchfield, D. W., Shilton, B. H., & Lüscher, B. (2008). Substrate-Assisted Catalysis by PARP10 Limits Its Activity to Mono-ADP- Ribosylation. Molecular Cell, 32(1), 57–69. 10.1016/j.molcel.2008.08.009

Kothe, G. O., Kitamura, M., Masutani, M., Selker, E. U., & Inoue, H. (2010). PARP is involved in replicative aging in Neurospora crassa. Fungal Genetics and Biology, 47(4), 297–309. 10.1016/j.fgb.2009.12.012

Kurtz, S., Phillippy, A., Delcher, A. L., Smoot, M., Shumway, M., Antonescu, C., & Salzberg, S. L. (2004). Versatile and open software for comparing large genomes. Genome Biology, 5(2), R12. 10.1186/gb-2004-5-2-r12

Langelier, M.-F., Eisemann, T., Riccio, A. A., & Pascal, J. M. (2018). PARP family enzymes: regulation and catalysis of the poly(ADP-ribose) posttranslational modification. Current Opinion in Structural Biology, 53, 187–198. 10.1016/j.sbi.2018.11.002

Letunic, I., & Bork, P. (2021). Interactive Tree Of Life (iTOL) v5: an online tool for phylogenetic tree display and annotation. Nucleic Acids Research, 49(W1), W293–W296. 10.1093/nar/gkab301

Leung, C. C. Y., & Glover, J. N. M. (2011). BRCT domains: easy as one, two, three. Cell Cycle (Georgetown, Tex.), 10(15), 2461–2470. 10.4161/cc.10.15.16312

Li, Y., Steenwyk, J. L., Chang, Y., Wang, Y., James, T. Y., Stajich, J. E., Spatafora, J. W., Groenewald, M., Dunn, C. W., Hittinger, C. T., Shen, X.-X., & Rokas, A. (2021). A genome-scale phylogeny of the kingdom Fungi. Current Biology, 31(8), 1653–1665.e5. 10.1016/j.cub.2021.01.074

Ma, L.-J., van der Does, H. C., Borkovich, K. A., Coleman, J. J., Daboussi, M.-J., Di Pietro, A., Dufresne, M., Freitag, M., Grabherr, M., Henrissat, B., Houterman, P. M., Kang, S., Shim, W.- B., Woloshuk, C., Xie, X., Xu, J.-R., Antoniw, J., Baker, S. E., Bluhm, B. H., … Rep, M. (2010). Comparative genomics reveals mobile pathogenicity chromosomes in Fusarium. Nature, 464(7287), 367–373. 10.1038/nature08850

Meyer-Ficca, M. L., Meyer, R. G., Coyle, D. L., Jacobson, E. L., & Jacobson, M. K. (2004). Human poly(ADP-ribose) glycohydrolase is expressed in alternative splice variants yielding isoforms that localize to different cell compartments. Experimental Cell Research, 297(2), 521–532. 10.1016/j.yexcr.2004.03.050

Mikolčević, P., Hloušek-Kasun, A., Ahel, I., & Mikoč, A. (2021). ADP-ribosylation systems in bacteria and viruses. Computational and Structural Biotechnology Journal, 19, 2366–2383. 10.1016/j.csbj.2021.04.023

Milo, S., Misgav, R. H., Hazkani-Covo, E., & Covo, S. (2019). Limited DNA repair gene repertoire in Ascomycete yeast revealed by comparative genomics. Genome Biology and Evolution. 10.1093/gbe/evz242

Nguyen, L.-T., Schmidt, H. A., von Haeseler, A., & Minh, B. Q. (2015). IQ-TREE: A Fast and Effective Stochastic Algorithm for Estimating Maximum-Likelihood Phylogenies. Molecular Biology and Evolution, 32(1), 268–274. 10.1093/molbev/msu300

Nicaud, J. (2012). Yarrowia lipolytica. Yeast, 29(10), 409–418. 10.1002/yea.2921

Novikova, O. (2009). Chromodomains and LTR retrotransposons in plants. Communicative & Integrative Biology, 2(2), 158–162. 10.4161/cib.7702

Ocaña-Pallarès, E., Williams, T. A., López-Escardó, D., Arroyo, A. S., Pathmanathan, J. S., Bapteste, E., Tikhonenkov, D. V., Keeling, P. J., Szöllősi, G. J., & Ruiz-Trillo, I. (2022). Divergent genomic trajectories predate the origin of animals and fungi. Nature, 609(7928), 747–753. 10.1038/s41586-022-05110-4

Otto, H., Reche, P. A., Bazan, F., Dittmar, K., Haag, F., & Koch-Nolte, F. (2005). In silico characterization of the family of PARP-like poly(ADP-ribosyl)transferases (pARTs). BMC Genomics, 6(1), 139. 10.1186/1471-2164-6-139

Perina, D., Mikoč, A., Ahel, J., Ćetković, H., Žaja, R., & Ahel, I. (2014). Distribution of protein poly(ADP-ribosyl)ation systems across all domains of life. DNA Repair, 23, 4–16. 10.1016/j.dnarep.2014.05.003

Pfaffl, M. W. (2001). A new mathematical model for relative quantification in real-time RT-PCR. Nucleic Acids Research, 29(9), 45e–445. 10.1093/nar/29.9.e45

Ray Chaudhuri, A., & Nussenzweig, A. (2017). The multifaceted roles of PARP1 in DNA repair and chromatin remodelling. Nature Reviews Molecular Cell Biology, 18(10), 610–621. 10.1038/nrm.2017.53

Rissel, D., & Peiter, E. (2019). Poly(ADP-Ribose) Polymerases in Plants and Their Human Counterparts: Parallels and Peculiarities. International Journal of Molecular Sciences, 20(7), 1638. 10.3390/ijms20071638

Rokas, A. (2022). Evolution of the human pathogenic lifestyle in fungi. Nature Microbiology, 7(5), 607–619. 10.1038/s41564-022-01112-0

Sauters, T. J. C., & Rokas, A. (2025). Patterns and mechanisms of fungal genome plasticity. Current Biology, 35(11), R527–R544. 10.1016/j.cub.2025.04.003

Schoutteten, N., Yurkov, A., Leroux, O., Haelewaters, D., Van Der Straeten, D., Miettinen, O., Boekhout, T., Begerow, D., & Verbeken, A. (2023). Diversity of colacosome-interacting mycoparasites expands the understanding of the evolution and ecology of *Microbotryomycetes*. Studies in Mycology, 106(1), 41–94. 10.3114/sim.2022.106.02

Selmecki, A., Forche, A., & Berman, J. (2010). Genomic Plasticity of the Human Fungal Pathogen Candida albicans. Eukaryotic Cell, 9(7), 991–1008. 10.1128/EC.00060-10

Semighini, C. P., Savoldi, M., Goldman, G. H., & Harris, S. D. (2006). Functional Characterization of the Putative *Aspergillus nidulans* Poly(ADP-Ribose) Polymerase Homolog PrpA. Genetics, 173(1), 87–98. 10.1534/genetics.105.053199

Smith, B. A., Neal, C. L., Chetram, M., Vo, B., Mezencev, R., Hinton, C., & Odero-Marah, V. A. (2013). The phytoalexin camalexin mediates cytotoxicity towards aggressive prostate cancer cells via reactive oxygen species. Journal of Natural Medicines, 67(3), 607–618. 10.1007/s11418-012-0722-3

Sowa, S. T., Bosetti, C., Galera-Prat, A., Johnson, M. S., & Lehtiö, L. (2022). An Evolutionary Perspective on the Origin, Conservation and Binding Partner Acquisition of Tankyrases. Biomolecules, 12(11), 1688. 10.3390/biom12111688

Tatusov, R. L. (2000). The COG database: a tool for genome-scale analysis of protein functions and evolution. Nucleic Acids Research, 28(1), 33–36. 10.1093/nar/28.1.33

Tawfik, O. K. and D. S. (2010). Enzyme Promiscuity: A Mechanistic and Evolutionary Perspective. Annual Review of Biochemistry, 79(1), 471–505. 10.1146/annurev-biochem-030409-143718

Teichmann, S. A., Rison, S. C. G., Thornton, J. M., Riley, M., Gough, J., & Chothia, C. (2001). The evolution and structural anatomy of the small molecule metabolic pathways in Escherichia coli. Journal of Molecular Biology, 311(4), 693–708. 10.1006/jmbi.2001.4912

Thumuluri, V., Almagro Armenteros, J. J., Johansen, A. R., Nielsen, H., & Winther, O. (2022). DeepLoc 2.0: multi-label subcellular localization prediction using protein language models. Nucleic Acids Research, 50(W1), W228–W234. 10.1093/nar/gkac278

Vlaardingerbroek, I., Beerens, B., Schmidt, S. M., Cornelissen, B. J. C., & Rep, M. (2016). Dispensable chromosomes in *Fusarium oxysporum* f. sp. *lycopersici*. Molecular Plant Pathology, 17(9), 1455–1466. 10.1111/mpp.12440

Vyas, S., Chesarone-Cataldo, M., Todorova, T., Huang, Y.-H., & Chang, P. (2013). A systematic analysis of the PARP protein family identifies new functions critical for cell physiology. Nature Communications, 4(1), 2240. 10.1038/ncomms3240

Vyas, S., Matic, I., Uchima, L., Rood, J., Zaja, R., Hay, R. T., Ahel, I., & Chang, P. (2014). Family-wide analysis of poly(ADP-ribose) polymerase activity. Nature Communications, 5(1), 4426. 10.1038/ncomms5426

Wadi, L., & Reinke, A. W. (2020). Evolution of microsporidia: An extremely successful group of eukaryotic intracellular parasites. PLOS Pathogens, 16(2), e1008276. 10.1371/journal.ppat.1008276

Wang, B., Yu, H., Jia, Y., Dong, Q., Steinberg, C., Alabouvette, C., Edel-Hermann, V., Kistler, H. C., Ye, K., Ma, L.-J., & Guo, L. (2020). Chromosome-Scale Genome Assembly of *Fusarium oxysporum* Strain Fo47, a Fungal Endophyte and Biocontrol Agent. Molecular Plant-Microbe Interactions®, 33(9), 1108–1111. 10.1094/MPMI-05-20-0116-A

Wang, J., Gao, Y., Xiong, X., Yan, Y., Lou, J., Guo, M., Noman, M., Li, D., & Song, F. (2025). Poly(ADP- ribose) polymerase FonPARP1-catalyzed PARylation of protein disulfide isomerase FonPdi1 regulates pathogenicity of Fusarium oxysporum f. sp. niveum on watermelon. International Journal of Biological Macromolecules, 291, 139046. 10.1016/j.ijbiomac.2024.139046

Wang, J., Gao, Y., Xiong, X., Yan, Y., Lou, J., Noman, M., Li, D., & Song, F. (2024). The Ser/Thr protein kinase FonKin4-poly(ADP-ribose) polymerase FonPARP1 phosphorylation cascade is required for the pathogenicity of watermelon fusarium wilt fungus Fusarium oxysporum f. sp. niveum. Frontiers in Microbiology, 15. 10.3389/fmicb.2024.1397688

Wu, B., Hao, W., & Cox, M. P. (2022). Reconstruction of gene innovation associated with major evolutionary transitions in the kingdom Fungi. BMC Biology, 20(1), 144. 10.1186/s12915-022-01346-8

Yang, H., Yu, H., & Ma, L.-J. (2020). Accessory Chromosomes in *Fusarium oxysporum*. Phytopathology®, 110(9), 1488–1496. 10.1094/PHYTO-03-20-0069-IA

Yong Zhang. (2019). EVOLUTION OF THE PATHOGENIC FUSARIUM OXYSPORUM THROUGH THE LENS OF COMPARATIVE GENOMICS [Doctor of Philosophy]. University of Massachusetts Amherst.

Yu, H., Yang, H., Haridas, S., Hayes, R. D., Lynch, H., Andersen, S., Newman, M., Li, G., Martínez-Soto, D., Milo-Cochavi, S., Hazal Ayhan, D., Zhang, Y., Grigoriev, I. V., & Ma, L.-J. (2023). Conservation and Expansion of Transcriptional Factor Repertoire in the Fusarium oxysporum Species Complex. Journal of Fungi, 9(3), 359. 10.3390/jof9030359

Zhang, Y., Yang, H., Turra, D., Zhou, S., Ayhan, D. H., DeIulio, G. A., Guo, L., Broz, K., Wiederhold, N., Coleman, J. J., Donnell, K. O., Youngster, I., McAdam, A. J., Savinov, S., Shea, T., Young, S., Zeng, Q., Rep, M., Pearlman, E., … Ma, L.-J. (2020). The genome of opportunistic fungal pathogen Fusarium oxysporum carries a unique set of lineage-specific chromosomes. Communications Biology, 3(1), 50. 10.1038/s42003-020-0770-2

