## Supplemental Table 5 for "The Ancient Origin and Dynamic Diversification of the Fungal Poly(ADP)-ribose Polymerase Protein Family"

Table S5. Human PARP Motif Reference

| **hPARP** | **Catalytic Motif** | **Activity** |
| --- | --- | --- |
| PARP1 | HYE | PAR |
| PARP2 | HYE | PAR |
| PARP3 | HYE | MAR/PAR |
| PARP4 | HYE | MAR |
| PARP5a | HYE | PAR |
| PARP5b | HYE | PAR |
| PARP6 | HYI | MAR |
| PARP7 | HYI | MAR |
| PARP8 | HYI | MAR |
| PARP9 | QYT | Inactive |
| PARP10 | HYI | MAR |
| PARP11 | HYI | MAR |
| PARP12 | HYI | MAR |
| PARP13 | YYV | Inactive |
| PARP14 | HYL | MAR |
| PARP15 | HYL | MAR |
| PARP16 | HYY | MAR |

List of hPARP proteins and their catalytic amino acids were adapted from (Vyas et al., 2014).
