## Supplemental Figure 1 for "The Ancient Origin and Dynamic Diversification of the Fungal Poly(ADP)-ribose Polymerase Protein Family"

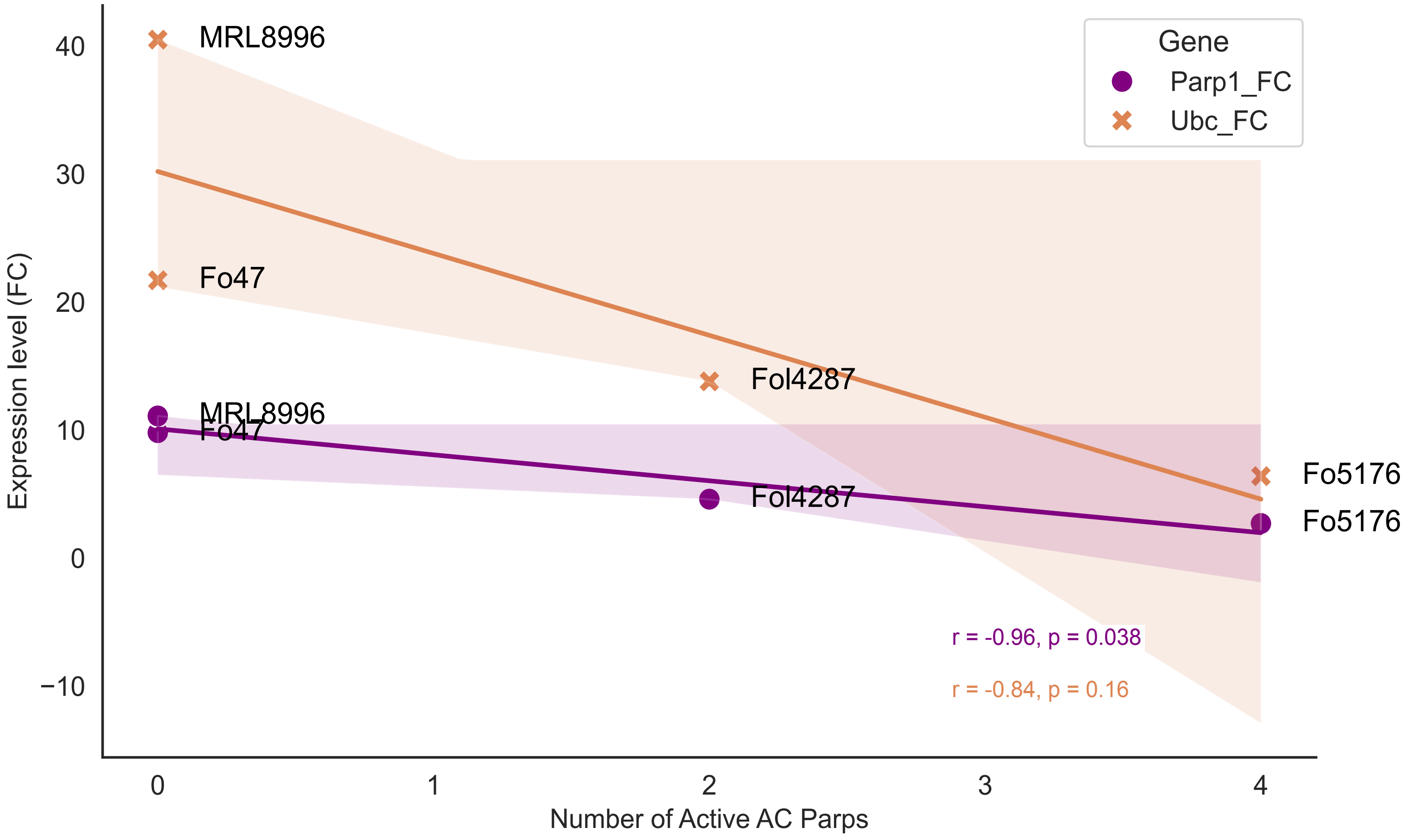


**Figure S1. Inverse relationship between core PARP expression and transcriptionally active accessory PARP genes.**
Scatter plots showing the relationship between the number of transcriptionally active accessory PARP genes induced following MMS treatment and the expression (fold change) of the core Parp1 (purple circles) and Parp-Ubc (orange crosses) genes across four F. oxysporum strains. Solid lines represent linear regression fits with shaded 95% confidence intervals. Pearson correlation coefficients (r) and corresponding p-values are indicated. Parp1 expression was negatively correlated with the number of transcriptionally active accessory PARP genes (r = -0.96, P = 0.038), whereas the correlation for Parp-Ubc expression was negative but not statistically significant (r = -0.84, P = 0.16).
